# *Ex vivo* detection of SARS-CoV-2-specific CD8+ T cells: rapid induction, prolonged contraction, and formation of functional memory

**DOI:** 10.1101/2020.08.13.249433

**Authors:** Isabel Schulien, Janine Kemming, Valerie Oberhardt, Katharina Wild, Lea M. Seidel, Saskia Killmer, Sagar, Franziska Daul, Marilyn Salvat Lago, Annegrit Decker, Hendrik Luxenburger, Benedikt Binder, Dominik Bettinger, Oezlem Sogukpinar, Siegbert Rieg, Marcus Panning, Daniela Huzly, Martin Schwemmle, Georg Kochs, Cornelius F. Waller, Alexandra Nieters, Daniel Duerschmied, Florian Emmerich, Henrik Mei, Axel Schulz, Sian Llewellyn-Lacey, David A. Price, Tobias Boettler, Bertram Bengsch, Robert Thimme, Maike Hofmann, Christoph Neumann-Haefelin

## Abstract

CD8+ T cells are critical for the elimination and long-lasting protection of many viral infections, but their role in the current SARS-CoV-2 pandemic is unclear. Emerging data indicates that SARS-CoV-2-specific CD8+ T cells are detectable in the majority of individuals recovering from SARS-CoV-2 infection. However, optimal virus-specific epitopes, the role of pre-existing heterologous immunity as well as their kinetics and differentiation program during disease control have not been defined in detail. Here, we show that both pre-existing and newly induced SARS-CoV-2-specific CD8+ T-cell responses are potentially important determinants of immune protection in mild SARS-CoV-2 infection. In particular, our results can be summarized as follows: First, immunodominant SARS-CoV-2-specific CD8+ T-cell epitopes are targeted in the majority of individuals with convalescent SARS-CoV-2 infection. Second, MHC class I tetramer analyses revealed the emergence of phenotypically diverse and functionally competent pre-existing and newly induced SARS-CoV-2-specific memory CD8+ T cells that showed similar characteristics compared to influenza-specific CD8+ T cells. Third, SARS-CoV-2-specific CD8+ T-cell responses are more robustly detectable than antibodies against the SARS-CoV-2-spike protein. This was confirmed in a longitudinal analysis of acute-resolving infection that demonstrated rapid induction of the SARS-CoV-2-specific CD8+ T cells within a week followed by a prolonged contraction phase that outlasted the waning humoral immune response indicating that CD8+ T-cell responses might serve as a more precise correlate of antiviral immunity than antibody measurements after convalescence. Collectively, these data provide new insights into the fine specificity, heterogeneity, and dynamics of SARS-CoV-2-specific memory CD8+ T cells, potentially informing the rational development of a protective vaccine against SARS-CoV-2.

## Introduction

Infections with the newly emerging coronavirus – severe acute respiratory syndrome coronavirus-2 (SARS-CoV-2) – cause the global outbreak of coronavirus disease 2019 (COVID-19)^1^. First cases occurred in December 2019 and as of mid-August 2020, roughly 20.3 million cases and 740.000 deaths have been documented. The clinical course of SARS-CoV-2 infections is highly variable and ranges from asymptomatic infections over mild courses with fever and cough to severe pneumonia and acute respiratory distress syndrome^2^. Identification of the determinants of immune protection is a prerequisite for the development of vaccines and therapeutic interventions.

Early data have indicated that SARS-CoV-2-specific CD8+ T cells are detectable in up to 70% of convalescent individuals targeting different viral proteins^3, 4, 5, 6, 7, 8^. However, these studies did not define individual immunodominant SARS-CoV-2-specific CD8+ T-cell epitopes, a pre-requisite for the *ex vivo* characterization of SARS-CoV-2-specific CD8+ T cells. Interestingly, in 20-50% of unexposed individuals, CD8+ T cells responding to SARS-CoV-2 peptide pools have also been observed^3, 5, 7, 9, 10^ indicating pre-existing virus-specific CD8+ T-cell response most likely due to exposure to “common cold” coronaviruses. All in all, currently, very little information is available about the abundance, phenotype, functional capacity and fate of pre-existing and newly induced SARS-CoV-2-specific CD8+ T-cell responses during the natural course of SARS-CoV-2 infection.

In this study, we therefore performed a high-resolution analysis of SARS-CoV-2-specific CD8+ T-cell responses by defining a set of novel optimal immunodominant SARS-CoV-2-specific CD8+ T-cell epitopes enabling *ex vivo* comparison of pre-existing and newly induced SARS-CoV-2-specific CD8+ T cells applying peptide-loaded MHCI-tetramer technology. By these analyses, we observed a rapid induction, prolonged contraction and versatile emergence of heterogeneous and functionally competent pre-existing and newly induced memory CD8+ T-cell responses in individuals with a mild course of SARS-CoV-2 infection that were more robust compared to the accompanied SARS-CoV-2 antibody response targeting spike that are frequently used to monitor SARS-CoV-2 infections.

## Results

### Definition of novel dominant virus-specific CD8+ T-cell epitopes in convalescent SARS-CoV-2-infected patients

We predicted SARS-CoV-2-derived 8-, 9- or 10-mer peptides with high affinity for 10 HLA class I alleles that are common in most populations world-wide (Extended Data Fig. 1). We selected 5 epitope candidate peptides for each of the following HLA alleles: A*01:01, A*02:01, A*03:01, A*11:01, A*24:02 as well as B*07:02, B*08:01, B*15:01, and B*40:01 and 8 epitope candidate peptides for B*44:02/03 (Table 1). In addition, we also included all 13 described SARS-CoV-1-specific CD8+ T cell epitopes that display 100% homology in SARS-CoV-2^8^ (Table 1). Next, we tested these 66 epitope peptides in 26 individuals with convalescent mild SARS-CoV-2 infection (Extended Data Table 1) in peptide-specific cell cultures. Importantly, we could detect SARS-CoV-2-specific CD8+ T-cell responses in 23/26 (88.4%) of the individuals, targeting a median of 4 epitopes (range 1-12) (Fig. 1A). Of note, the HLA-A*02:01-restricted epitopes that had been pre-described for SARS-CoV-1 and that are completely conserved in SARS-CoV-2 (Table 1) were only rarely targeted in our cohort (Fig. 1B, Table 1). However, 33/53 (62.3%) SARS-CoV-2-specific epitope candidates predicted in our study could be confirmed (Table 1, depicted in bold). The strongest responses were observed for epitopes A*01/ORF3a_207-215_, A*02/ORF3a_139-147_, and B*07/N_105-113_ with a median of 8.3%, 8.4%, and 62.6% of CD8+ T cells producing IFN-γ after peptide-specific culture, respectively (Fig. 1B/C). Taking the protein length into account, we observed a relative overrepresentation of nucleocapsid- and ORF3a-specific CD8+ T-cell responses (Fig. 1D). Despite the superior immunogenicity reflected by the relative overrepresentation of nucleocapsid and ORF3a, the absolute majority of detected responses (57/110 [51.2%]) targeted ORF1ab (Fig. 1D). The T-cell epitopes were restricted by both HLA types, HLA-A and HLA-B to a similar extend (Fig. 1E). Interestingly, we were able to detect SARS-CoV-2-specific CD8+ T-cell responses in all nine convalescent individuals that were seronegative for anti-SARS-CoV-2 S antibodies (n=6) or had borderline test results (n=3) (Fig. 1F). Next, we set out to determine whether the identified SARS-CoV-2-specific CD8+ T cell epitopes are unique to SARS-CoV-2-exposed individuals. For this analysis, we tested a cohort of 25 healthy volunteers with comparable characteristics regarding gender and age compared to our SARS-CoV-2 cohort. Blood samples were obtained before August 2019 and thus prior to a possible exposure to SARS-CoV-2 (Extended Data Table 1) and tested for the presence of virus-specific CD8+ T cells in the same way as the individuals with convalescent SARS-CoV-2 infection. We observed only very low virus-specific IFN-γ and TNF CD8+ T-cell responses in 6 individuals (5 individuals with a single response and 1 individual with 5 responses) (Fig.1G, Table 1, Extended Data Fig.2A) and TNF without IFN-γ responses in additional 4 individuals (Extended Data Fig.2A/C). The only epitope that was targeted by IFN-γ secreting CD8+ T cells in more than one SARS-CoV-2-naïve individual was epitope B*07/N_105-113_ (Extended Data Fig.2A). Of note, this is the SARS-CoV-2-specific epitope in our study with the highest conservation between SARS-CoV-2 and “common cold” corona viruses (Extended Data Fig.2B, Extended Data Table 2). In summary, these results reveal that the majority of identified SARS-CoV-2-specific CD8+ T-cell epitopes that were dominantly targeted in convalescent individuals with mild SARS-CoV-2 infection, show little evidence for cross-recognition in SARS-CoV-2-naïve individuals.

**Table 1:**
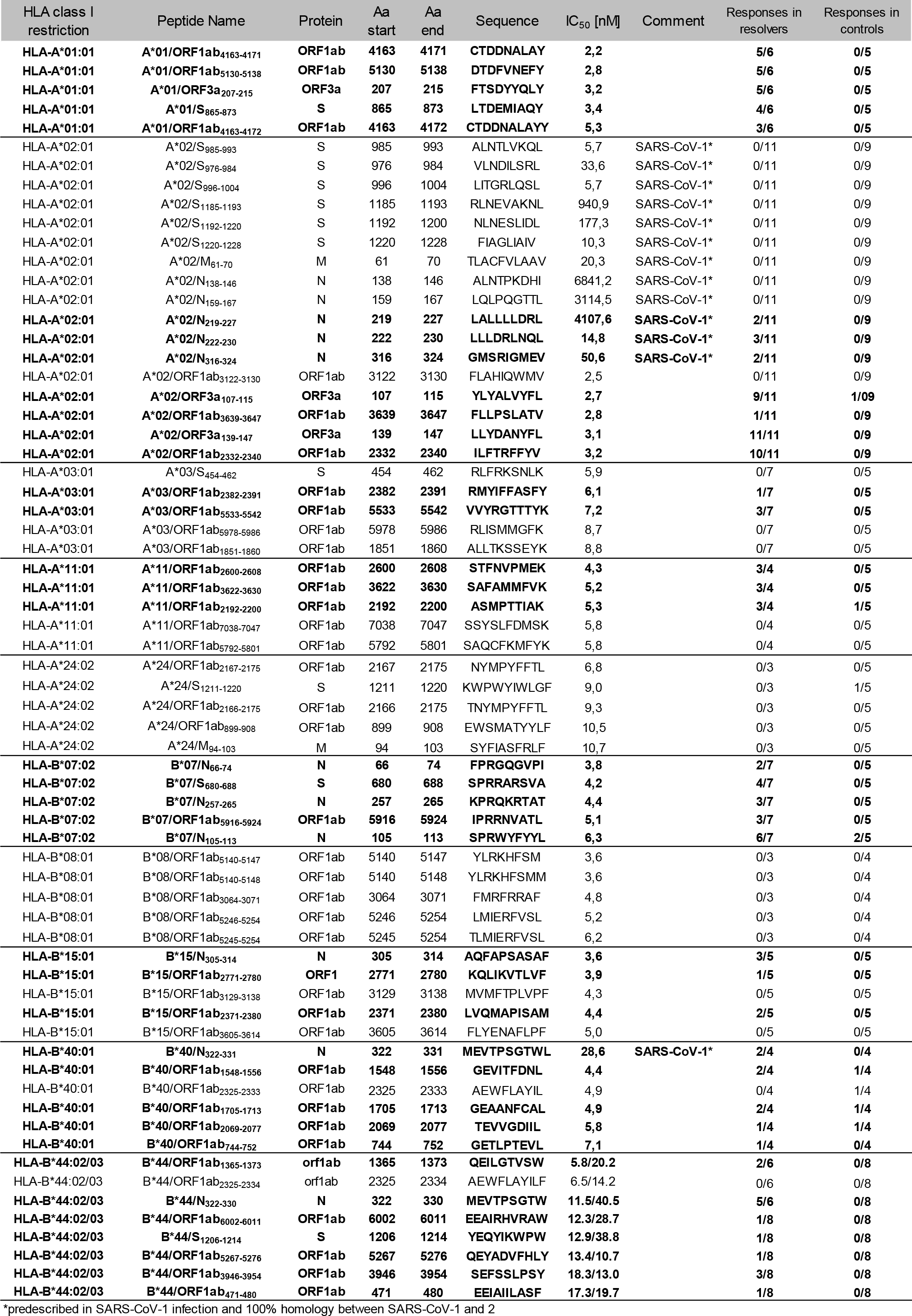
SARS-CoV-2-derived peptides used for analyses of SARS-CoV-2-specific CD8+ T cell responses. Peptides in bold indicate epitopes confirmed in this study.

**Figure 1:**
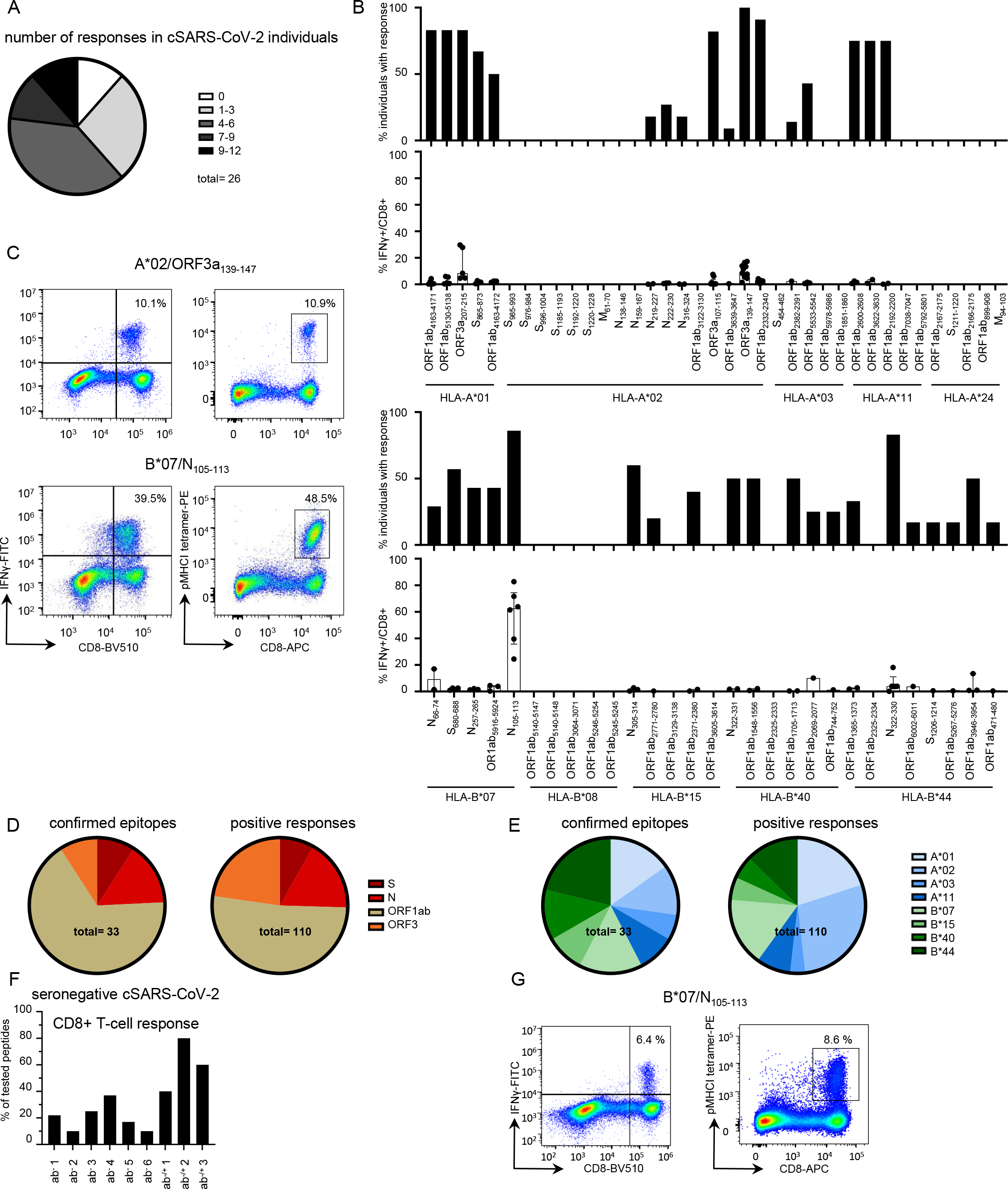
Definition of dominant SARS-CoV-2-specific CD8+ T-cell epitopes in convalescent SARS-CoV-2 individuals. **(A)** Pie chart illustrating the number of epitopes recognized per tested individual. **(B)** % of convalescent SARS-CoV-2 individuals with positive response towards HLA-A- and HLA-B-restricted SARS-CoV-2 peptides as well as the strength of individual responses as % IFN-γ+ of CD8+ T cells. **(C)** Representative dot plots showing pMHCI-tetramer stainings and IFNγ production of A*02/ORF3a_139-147_ and B*07/N_105-113_-specific CD8+ T cells after 14-days *in vitro* expansion. Numbers refer to the respective percentage of pMHCI-tetramer+ and IFN-γ+ cells among CD8+ T cells. Confirmed epitopes and total positive responses are depicted regarding their location within the SARS-CoV-2 genome **(D)** and according to their HLA restriction **(E). (F)** CD8+ T-cell responses in SARS-CoV-2 antibody seronegative or borderline positive convalescent patients as percentage of responses out of all peptides tested matching the patient’s HLA alleles. **(G)** Exemplary dot plots showing a pMHCI-tetramer staining and IFNγ production of HLA-B*07/N_105-113_-specific CD8+ T cells from a historic control after 14-days *in vitro* expansion. Numbers refer to the respective percentage of pMHCI-tetramer+ and IFN-γ+ cells among CD8+ T cells. Bar charts show the median with IQR.

### Phenotypic memory characteristics of ex vivo detectable HLA-A and HLA–B-restricted SARS-CoV-2-specific CD8+ T cells

To evaluate the phenotypic characteristics of SARS-CoV-2-specific memory CD8+ T-cell populations, by using a set of novel MHC I tetramers we analyzed *ex vivo* SARS-CoV-2-specific CD8+ T cells targeting six immunodominant epitopes (A*01/ORF3a_207-215_, A*01/ORF1ab_4163-4172_, A*02/ORF3a_139-147_, B*07/N_105-113_ B*44:03/N_322-330_, B*44:03/ORF1ab_3946-3954_) in comparison to influenza (FLU)-specific CD8+ T cells (A*02/Flu-M1_58-66_) in a cohort of 18 convalescent individuals following a mild course of infection. In order to increase the detection rate and to allow subsequent in-depth phenotypic analysis, we performed peptide-loaded MHC I tetramer-based enrichment (Fig. 2A). Remarkably, we could detect SARS-CoV-2-specific CD8+ T cells *ex vivo* in nearly all tested convalescent individuals (Fig. 2B). The *ex vivo* frequencies of SARS-CoV-2-specific CD8+ T cells targeting A*01/ORF3a_207-215_; A*01/ORF1ab_4163-4172_; A*02/ORF3a_139-147_; B*44:03/N_322-330_ and B*44:03/ORF1ab_3946-3954_ were similar (Fig. 2B). CD8+ T cells targeting B*07/N_105-113_ were present in slightly higher frequencies compared to other SARS-CoV-2-specific CD8+ T-cell populations reaching levels of A*02/Flu-M1_58-66_-specific CD8+ T cells (Fig. 2B). This probably reflects heterologous stimulation of pre-existing B*07/N_105-113_-specific CD8+ T cells (Extended Data Fig.2A). SARS-CoV-2-specific CD8+ T-cell populations in convalescent individuals were composed of naïve (T_naive_), central memory (T_CM_), effector memory 1 (T_EM1_), effector memory 2 (T_EM2_), effector memory 3 (T_EM3_) and terminally differentiated effector memory expressing RA (T_EMRA_) T-cell subsets irrespective of the targeted epitope (Extended Data Fig. 3A/B). The presence of an only minor T_naive_ subset fraction among all tested SARS-CoV-2-specific CD8+ T cells supports that these cells have been efficiently primed during the infection. In comparison to HLA-B-restricted SARS-CoV-2-specific CD8+ T cells, HLA-A restricted virus-specific CD8+ T cells showed a shift towards the early differentiated T_CM_ and T_EM1_ subset (Extended Data Fig. 3B). Similar results were obtained by applying the CX_3_CR1-based definition of memory T-cell subsets (Extended Data Fig. 3C). To more comprehensively compare the phenotypes of the different SARS-CoV-2-specific CD8+ T cells we performed t-distributed stochastic neighbor embedding (t-SNE) of all analyzed virus-specific CD8+ T cells from the tested convalescent individuals (Fig. 2C). Topographical clustering of SARS-CoV-2-specific CD8+ T cells separated these cells according to their HLA restriction (left panel) dominating the respective differences associated with the targeted viral proteins (right panel). This was further supported by multidimensional scaling (MDS) analysis (Fig. 2C). HLA-A-restricted SARS-CoV-2-specific CD8+ T cells were characterized by a cluster of markers including CD38, PD-1 and TOX that are associated with antigen recognition as well as CD28 and TCF-1 labelling less differentiated cells (Fig. 2D). In contrast, HLA-B-restricted SARS-CoV-2-specific CD8+ T cells cluster based on CD45RA, CD57, KLRG1, CD25, CX_3_CR1 and high T-BET expression probably reflecting a more terminally differentiated effector cell state (Fig. 2D and Extended Data Fig. 3D). Of note, FLU-A*02/M1_58_-specific CD8+ T cells showed differences to HLA-A and B-restricted SARS-CoV-2-specific CD8+ T cells (Fig. 2C/D). In particular, FLU A*02/M1_58-66_-compared to SARS-CoV-2-specific CD8+ T cells expressed higher levels of CD127 and BCL2, both important factors for the homeostatic maintenance of memory T cells while the T-cell memory-associated transcription factors TCF-1 and FOXO1 were similarly expressed (Fig. 2E, Extended Data Fig. 4A). The reduced BCL-2 expression of SARS-CoV-2-specific CD8+ T cells was most prominent among the early differentiated T_CM_ and T_EM1_ subsets that have the highest BCL-2 expression among memory T-cell subsets in general (Extended Data Fig. 4B). Importantly, BCL-2 expression of SARS-CoV-2-specific CD8+ T cells correlated with the days after onset of symptoms (Fig. 2F). Thus, SARS-CoV-2-specific CD8+ T cells are most probably within the dynamic process of establishing a *bona fide* long-lasting memory compartment. In summary, circulating SARS-CoV-2-specific CD8+ T cells are frequently detectable *ex vivo* in convalescent individuals and are composed of different *bona fide* memory subsets with an additional layer of phenotypic heterogeneity based on the HLA restriction.

**Figure 2:**
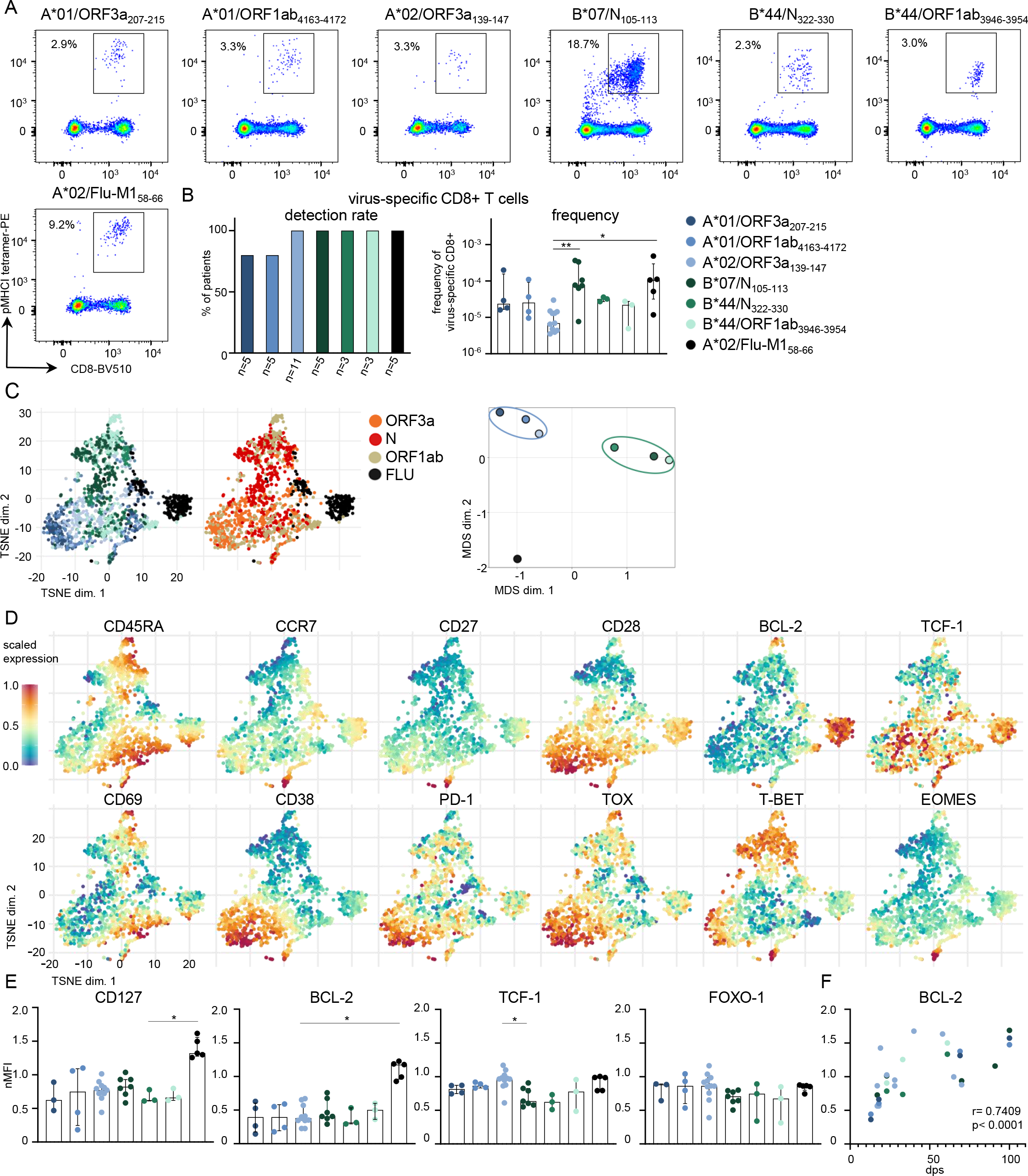
Phenotypic differences of *ex vivo* detectable virus-specific CD8+ T cells in SARS-CoV-2 convalescent individuals. **(A)** Representative dot plots showing A*01/ORF3a_207-215_-, A*01/ORF1ab_4163-4172_-, A*02/ORF3a_139-147_-, B*07/N_105-113_-, B*44/N_322-330_-, B*44/ORF1ab_3946-3954_- and A*02/Flu-M_158-66_-, specific CD8+ T cells ex vivo after pMHCI tetramer-based enrichment. **(B)** The detection rate (left) and frequency (right) of epitope-specific CD8+ T cells was determined. **(C)** t-distributed stochastic neighbor embedding (t-SNE) representation of flow cytometry data, which were derived from 18 SARS-CoV-2 convalescent individuals, comparing SARS-CoV-2-specific CD8+ T cells by their HLA restriction (left) and by their targeted viral proteins (middle). Multidimensional scaling (MDS) analysis comparing the similarity of HLA-A and HLA-B-restricted SARS-CoV-2-specific epitopes (right). (**D**) Expression levels of CD45RA, CCR7, CD27, CD28, BCL-2, TCF-1, CD69, CD38, PD-1, TOX, T-BET and EOMES are plotted on the t-SNE plot. Expression levels are color-coded: blue, low expression; red, high expression. (**E**) Mean fluorescence intensity of CD127, BCL-2, TCF-1 and FOXO-1 of virus-specific CD8+ T cells normalized to mean fluorescence intensity of naïve CD8+ T cells (nMFI). (**F**) Correlation of BCL-2 expression with date post symptom onset (dps). Bar charts show the median with IQR. Statistical significance was assessed by Kruskal-Wallis test including Dunn’s multiple comparisons test and Spearman correlation. (*P<0.05; **P<0.01; ***P<0.001; ****P<0.0001)

### Similar vigorous functional capacity of pre-existing and newly induced SARS-CoV-2-specific memory T cells

Next, we assessed the functional capacity of SARS-CoV-2-specific compared to FLU-specific memory CD8+ T cells *in vitro* (Fig. 3A). As shown in Fig. 3B after two weeks of *in vitro* expansion, we detected comparable frequencies of SARS-CoV-2 B*07/N_105-113_- and FLU A*02/M1_58-66_-specific CD8+ T cells that were higher compared to the other tested SARS-CoV-2-specific CD8+ T cells (left panel). However, when analyzing the expansion index, a measure taking the input number of virus-specific CD8+ T cells into account, we observed comparable *in vitro* expansion capacities of the analyzed SARS-CoV-2- and FLU-specific CD8+ T cells (Fig. 3B, right panel). Thus, the increased frequencies of SARS-CoV-2 B*07/N_105_-specific CD8+ T cells after peptide-specific CD8+ T-cell expansion most probably reflect a higher *ex vivo* frequency of these cells. We also analyzed cytokine production (IFN-γ and TNF) and degranulation as indicated by CD107a expression in relation to the frequency of virus-specific CD8+ T cells after expansion in order to have an approximation for the effector functions of SARS-CoV-2-specific CD8+ T cells. As shown in Fig. 3C, irrespective of the targeted epitope, the cytokine production and degranulation capacity of SARS-CoV-2-specific CD8+ T cells is similar to A*02/Flu-M1_58-66_-specific CD8+ T cells. In a next set of experiments, we addressed the question whether SARS-CoV-2 B*07/N105-113-specific memory CD8+ T-cell responses observed in SARS-CoV-2 convalescent individuals differ compared to “common cold” corona viruses-exposed individuals. For this, we analyzed SARS-CoV-2 B*07/N_105-113_-specific CD8+ T cells in historical samples (banked before August 2019) of six B*07:02 positive individuals (Fig. 3D). We detected SARS-CoV-2 B*07/N_105-113_-specific CD8+ T cells *ex vivo* in three out of six historic controls, however, at lower frequencies compared to SARS-CoV-2 convalescent individuals (Fig. 3D) indicating a heterologous boost expansion in the latter cohort. As depicted in Fig. 3E, the CD45RA/CCR7/CD27-based T-cell subset distribution revealed a slight shift towards the further differentiated T_EM3_ subset in SARS-CoV-2 convalescent individuals again supporting heterologous stimulation. However, we did not observe differences in expansion and cytokine production of SARS-CoV-2 B*07/N105-113-specific CD8+ T-cell population in SARS-CoV-2 convalescent individuals compared to historic controls (Fig. 3F). Taken together, these observations suggest that pre-existing and newly induced SARS-CoV-2-specific CD8+ T cells establish a functionally competent *bona fide* memory response similar to FLU-specific CD8+ T cells.

**Figure 3:**
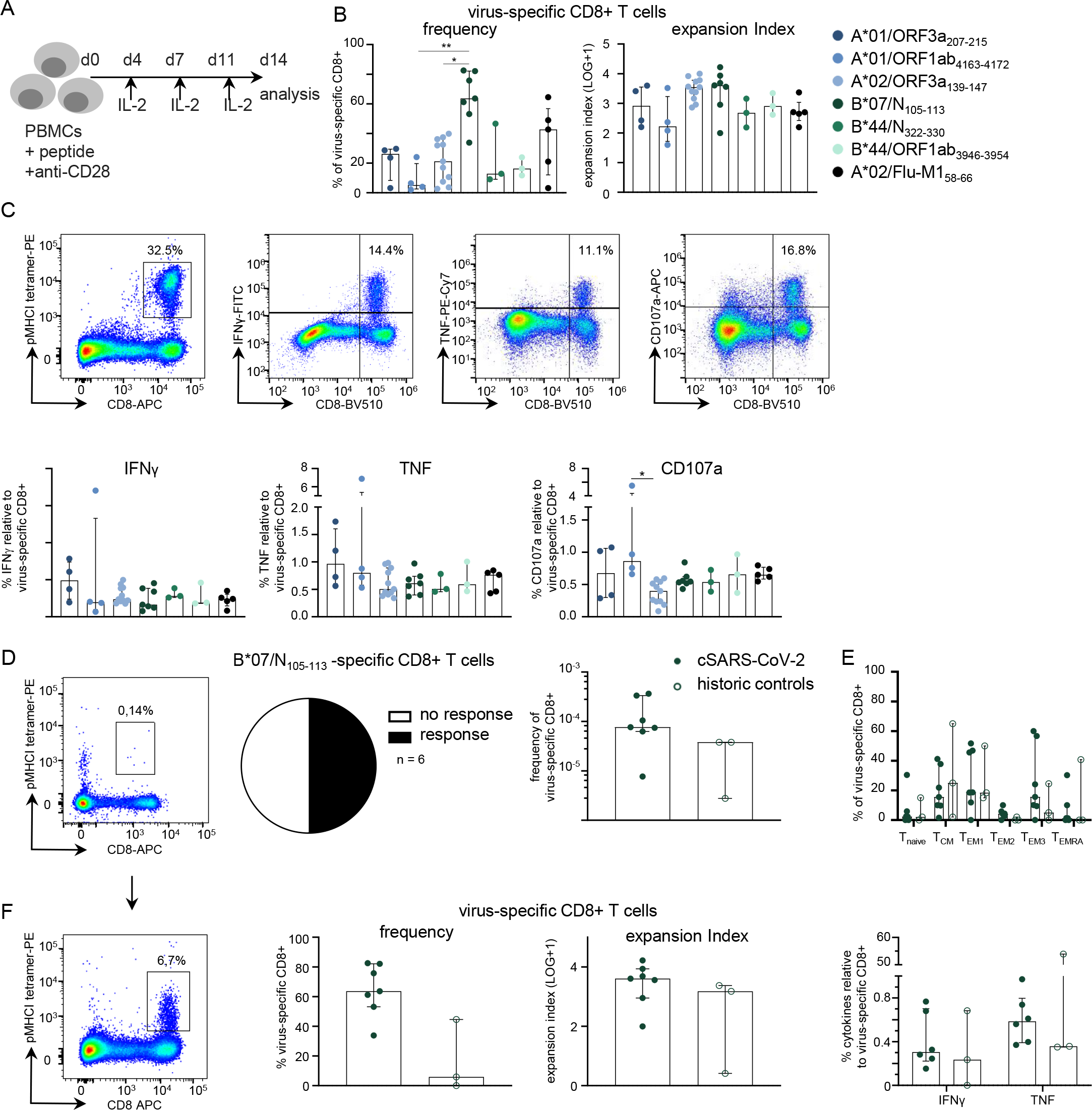
Similar vigorous functional capacity of pre-existing and newly induced SARS-CoV-2-specific memory CD8+ T cells. **(A)** Workflow illustrating the experimental set-up for the peptide-specific *in vitro* expansion of CD8+ T cells. 1.5×10^6^ PBMCs were stimulated with SARS-CoV-2-specific peptides and anti-CD28 mAb and expanded for 14 days in the presence of IL-2. **(B)** After 14 days of in vitro expansion the % of virus-specific CD8+ T cells (left) and the expansion index (right) of the respective epitope-specific CD8+ T cells were calculated. **(C)** Representative dot plots showing SARS-CoV-2-specific CD8+ T cells, as well as IFN-γ-, TNF and CD107a-producing CD8+ T cells after 14-days *in vitro* expansion (top). The percentage of IFNγ-, TNF- and CD107a-producing CD8+ T cells in relation to the frequency of epitope-specific CD8+ T cells was determined after 14-days *in vitro* expansion (bottom). **(D)** Representative dot plot showing virus-specific CD8+ T cells ex vivo after B*07/N_105-113_ tetramer-based enrichment (left), pie chart depicting the number of positive responses of patients tested (middle) and frequency of B*07/N_105-113_-specific CD8+ T cells in historic controls in comparison to convalescent SARS-CoV-2 individuals (cSARS-CoV-2) (right). **(E)** Distribution of T-cell memory subsets, Tnaive, TCM, TEM1, TEM2, TEM3 and TEMRA of B*07/N_105-113_-specific CD8+ T cells in historic controls compared to SARS-CoV2 convalescent individuals. **(F)** Representative dot plot showing virus-specific CD8+ T cells after 14 days in vitro expansion (left). The frequency and expansion index (middle) of virus-specific CD8+ T cells was determined. Expression of IFN-γ and TNF in percentage relative to the frequency of epitope-specific CD8+ T cells is shown. Bar charts show the median with IQR. Statistical significance was assessed by Kruskal-Wallis testing including Dunn’s multiple comparisons test. (*P<0.05; **P<0.01; ***P<0.001; ****P<0.0001)

### Rapid expansion and prolonged contraction of newly induced SARS-CoV-2-specific CD8+ T cells

We had the unique opportunity to longitudinally follow the SARS-CoV-2-specific CD8+ T-cell response before, during and after SARS-CoV-2 infection in an HLA-B*44:03^+^ individual with a defined infection event (Fig. 4A). As depicted in Fig. 4B and Extended Data Fig. 5A, SARS-CoV-2 B*44:03/N_322-330_- and SARS-CoV-2 B*44:03/ORF1ab_3946-3954_-specific CD8+ T cells were clearly expanded as early as 7 days post infection together with symptom onset. Importantly, both T-cell populations were not detectable prior to the SARS-CoV-2 infection clearly indicating novel priming (Fig. 4B and Extended Data Fig. 5A). The kinetics of both T cell responses were similar and the contraction phase lasted at least 70 days with SARS-CoV-2-specific CD8+ T cells still detectable at significant frequencies (approx. 1*10^-5^) 109 days post infection. Interestingly, the serum anti-SARS-CoV-2 S protein antibody titer fell below the upper detection limit at 84 days post infection (Fig. 4C) while the virus-specific CD8+ T cells remained detectable at this exact time point and also at later follow-up time points. Next, we performed deep profiling of SARS-CoV-2-specific CD8+ T cells including T-cell differentiation and activation markers, transcription factors, inhibitory receptors and pro-survival factors by using flow and mass cytometry to more comprehensively understand the T-cell phenotype and differentiation program during the course of infection. Diffusion map embedding combining flow cytometry data of SARS-CoV-2 B*44:03/N_322-330_- and SARS-CoV-2 B*44:03/ORF1ab_3946-3954_-specific CD8+ T cells indicated a continuous relationship between all SARS-CoV-2-specific CD8+ T cells longitudinally collected during and after infection, with cells from early time points after infection and those from late time points at opposing ends, reflecting a dynamic differentiation of the virus-specific CD8+ T-cell response (Fig. 4D, Extended Data Fig. 5B/C/D). SARS-CoV-2-specific CD8+ T cells collected at later time points compared to earlier time points post infection clustered more closely within the diffusion map suggesting a higher degree of similarity and likely establishment of a steady state at the memory phase of the T-cell response (Fig. 4D, Extended Data Fig. 4C). Based on the linearity of the differentiation program suggested by the diffusion map analysis, we performed single-cell trajectory detection using wanderlust analysis^17^ of CyTOF data to understand the differentiation trajectories in more detail (Extended Data Fig. 6A). This analysis showed that a small fraction of virus-specific T cells identified after one week of infection with a CD28+ TCF-1+ CD127+ CD45RA+ phenotype may represent the precursor population of the large pool of effector cells (Extended Data Fig. 6A). As indicated by these wanderlust (Extended Data Fig. 6A) and diffusion map (Fig. 4D) analyses, Phenotyping by Accelerated Refined Community-partitioning (PARC) of mass cytometry data confirmed a significant shift of SARS-CoV-2-specific CD8+ T cells from an early effector state characterized by a high expression of the activation markers CD38, CD39 or PD-1 together with Ki-67 towards a TEM differentiation program with high expression of CD45RA, CX_3_CR1, KLRG1 and CD57 with little involvement of T_CM_ cells (Fig. 4E, Extended Data Fig. 6B-D). These changes were also apparent on non-MHC I tetramer+ CD8+ T cells (Extended Data Fig. 6B/E) suggesting broad activation of virus-specific responses targeting other epitopes. Within a time-span of more than 100 days post infection, we did not detect major changes in the *in vitro* functional capacity (expansion, cytokine production and degranulation) of both SARS-CoV-2 B*44:03/N_322-330_- and SARS-CoV-2 B*44:03/ORF1ab_3946-3954_-specific CD8+ T-cell populations (Fig. 4F-H). Together, these findings suggest an ongoing efficient control of or protection from SARS-CoV-2 infection by virus-specific CD8+ T cells even at late time points, when antibodies may already have waned.

**Figure 4:**
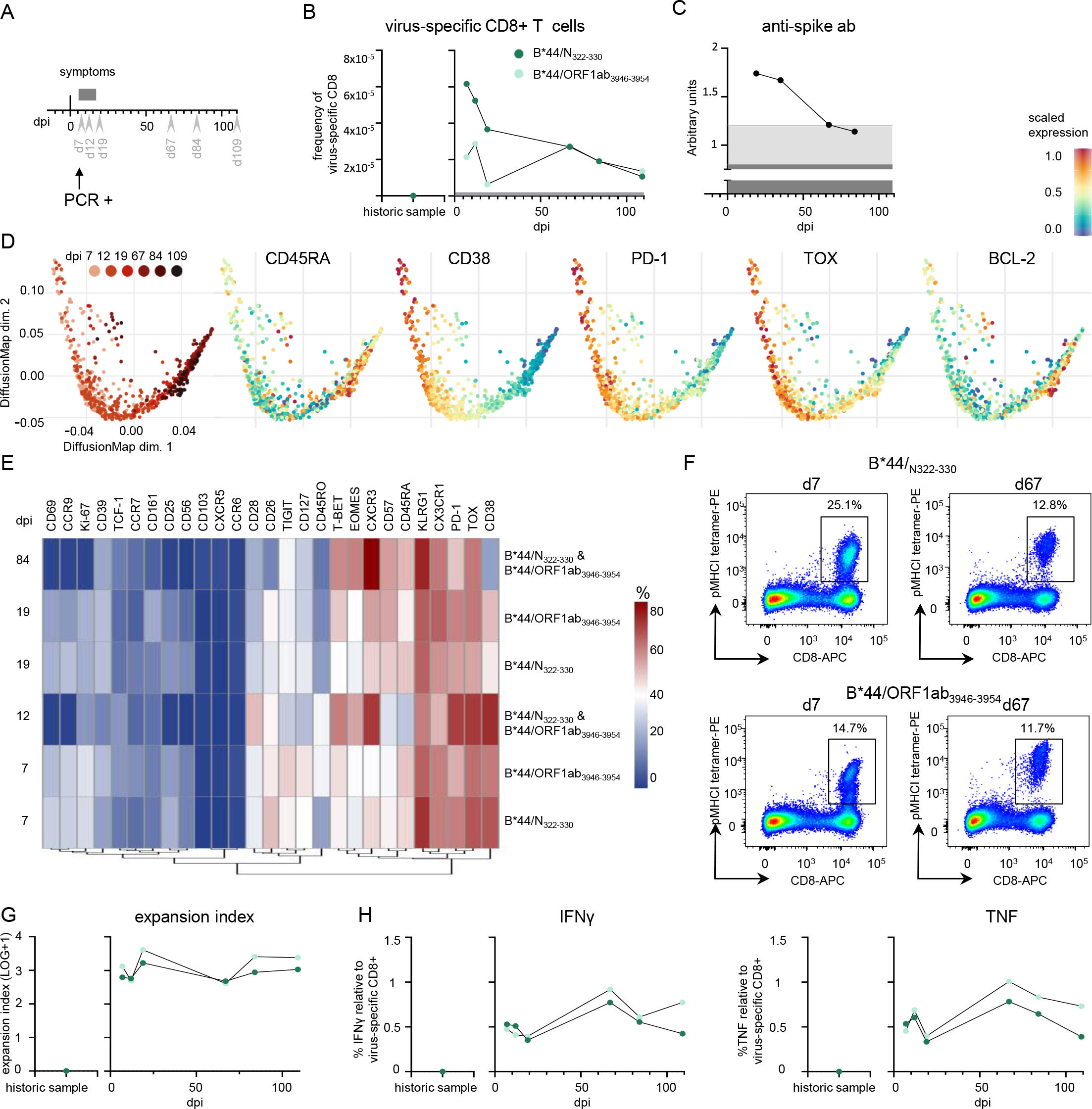
Rapid expansion and prolonged contraction of newly induced SARS-CoV-2-specific CD8+ T cells. **(A)** Timeline depicting the longitudinal sampling for the SARS-CoV-2-infected patient analyzed. Bleed dates (gray arrow heads), symptoms (dark grey bar) and positive PCR testing are shown at days post infection (dpi). **(B)** The frequency of B*44/N_322-330_- and B*44/ORF1ab_3946-3954_-specific T cells in the patient is indicated at dpi together with a historic sample. Grey line indicates detection threshold **(C)** Timeline depicting the anti-spike antibody response in arbitrary units at dpi. Light grey and dark grey background color indicate the area bellow the upper and lower detection limit, respectively **(D)** Diffusion map representation of flow cytometry data, which were derived from longitudinal analysis, demonstrating the diffusion of B*44/N_322-330_- and B*44/ORF1ab_3946-3954_-specific T cells in relation to dpi which is distinguished by a color gradient from light (early time points) to dark red (late time points) color. Protein expression levels are plotted on the diffusion map. **(E)** The dynamic expression profile of SARS-CoV-2-specific CD8+ T cells is visualized in a heatmap. Heatmap coloring represents % of virus-specific CD8+ T cells expressing a given marker; blue, low expression; red, high expression. **(F)** Dot plots showing pMHCI-tetramer stainings of B*44/N_322-330_- and B*44/ORF1ab_3946-3954_.specific CD8+ T cells after 14-days *in vitro* expansion at different time points post infection. Numbers refer to the respective percentage of pMHCI-tetramer+ cells among CD8+ T cells. Expansion index of virus-specific CD8+ T cells **(G)** and expression of IFNγ and TNF **(H)** in percentage relative to the frequency of epitope-specific CD8+ T cells after 14-days *in vitro* expansion at dpi and historic sample.

## Discussion

Here, we have defined a set of immunodominant CD8+ T-cell epitopes that were targeted in the majority of tested convalescent individuals of a Caucasian cohort after a mild course of SARS-CoV-2 infection. This even exceeds the previously reported high detection rate of T-cell responses in up to 70% of convalescent individuals in different cohorts using peptide pools for T-cell stimulation^3, 4, 5, 6, 7^, most likely owing to our more specific approach. These analyzed cohorts of convalescent individuals comprised citizen of the UK^7^, Sweden^4^, Singapore^5^ and California/USA^3^. Thus, a SARS-CoV-2-specific CD8+ T-cell memory is robustly induced on a global population level. This observation gains further relevance when taking into account that we could detect SARS-CoV-2-specific CD8+ T cells in individuals being seronegative for anti-SARS-CoV-2 spike antibodies indicating a higher sensitivity for detecting SARS-CoV-2-specific T cells compared to antibodies to prove a recent SARS-CoV-2 infection. A faster waning of the antibody response is also suggested by our longitudinal analysis of an individual with a mild disease course before, during and after SARS-CoV-2 infection where T-cell but not antibody responses were still detectable long after clinical resolution of infection. In addition, this is in line with previous reports in the context of SARS-CoV-1 infection reporting long-lasting detection of virus-specific CD8+ T cells in contrast to antibodies^18, 19^. Altogether, our findings indicate that SARS-CoV-2-specific CD8+ T cells represent a major determinant of immune protection on an individual as well as population level.

The set of CD8+ T-cell epitopes defined in this study is mainly composed of epitopes that have not been described to date and for which there is in most cases no evidence for pre-existing immunity as assessed by testing historic control samples collected before August 2019. The individuals included into our historic control cohort had no history of SARS-CoV-1 exposure that potentially impacts the virus-specific CD8+ T-cell response towards SARS-CoV-2 due to high sequence homology within T-cell epitopes^8^ and long-living cross-reactive virus-specific CD8+ T cells^5, 19^. The here-identified SARS-CoV-2-specific CD8+ T-cell responses target different structural and non-structural proteins with a specific focus against ORF1ab, in agreement with its protein length. However, taking the protein length into account, we observed a relative dominance of the N protein and ORF3a as targets. In line with Grifoni *et al.*, this finding emphasizes the broad recognition of SARS-CoV-2 by virus-specific CD8+ T cells and extends other previous studies restricted to structural proteins^3, 4, 6^ and omitting ORF1ab^7^. Thus, the SARS-CoV-2-specific CD8+ T-cell response interferes with different stages of the viral lifecycle, e.g. by targeting ORF1ab products necessary for the early viral replication and transcription^20^. Clearly, our approach to define optimal CD8+ T-cell epitopes based on a *in silico* prediction has the limitation that it does not completely cover the entire viral genome as it is the case in studies that have used overlapping peptides^3, 7, 9^. However, an advantage of this approach is the definition of exact, single and optimal CD8+ T-cell epitopes including HLA restriction. With this, our data revealed that there is no clear dominance of HLA-A or B-restricted epitopes that are targeted by SARS-CoV-2-specific CD8+ T cells indicating an evenly broad and robust induction of an antiviral CD8+ T-cell response among individuals.

Importantly, the definition of optimal epitopes also allowed a comparative *ex vivo* detection and characterization of SARS-CoV-2-specific CD8+ T cells after peptide-loaded MHC I tetramer-based enrichment. The highest frequency *ex vivo* was detectable for B*07/N_105-113_-specific CD8+ T cells which was also in agreement with their strong peptide-specific expansion. Indeed, B*07/N_105-113_-specific CD8+ T cells were similarly frequent as A*02/Flu-M1_58-66_-specific CD8+ T cells. Since the sequence homology of the B*07/N_105-113_ epitope among the corona viruses including “common cold” corona viruses is high and since we also identified pre-existing B*07/N_105-113_-specific CD8+ T cells in historic controls, the higher frequency of these CD8+ T cells in SARS-CoV-2 convalescent individuals most probably reflects heterologous boosting. Of note, in the study by Peng *et al*.^7^, this epitope was also included in the dominant overlapping peptide pool showing CD8+ T-cell responses in 11 out of 42 individuals. Interestingly, 10 out of theses 11 individuals expressed HLA-B*07:02. However, we did not observe clear phenotypical and functional differences between the potentially pre-existing B*07/N_105-113_-specific CD8+ T cells and other newly induced SARS-CoV-2-specific CD8+ T cells either targeting structural or non-structural proteins. One reason for this may be a rapid and strong induction also of newly induced SARS-CoV-2-specific CD8+ T cells that we have detected in one individual followed longitudinally during mild SARS-CoV-2 infection. Furthermore, the phenotypic and functional characteristics of SARS-CoV-2-specific CD8+ T cells were quite similar to the immunodominant A*02/Flu-M158-66-specific CD8+ T cells representing classical, fully functional memory T cells^21^. The examined lower BCL-2 expression of SARS-CoV-2-compared to FLU-specific CD8+ T cells is most probably due to the fact that the SARS-CoV-2-specific CD8+ T-cell memory response was not in a resting steady state. This hypothesis is corroborated by our findings that BCL-2 expression increased with time after SARS-CoV-2 infection and that the contraction phase of SARS-CoV-2-specific CD8+ T cells was apparently prolonged in the individual that we longitudinally followed during SARS-CoV-2 infection. Taken together, these comprehensive analyses revealed that a fully functional immune response is generated by both, pre-existing and newly induced virus-specific CD8+ T cells, irrespective of the targeted viral protein and the HLA restriction. The established functionally competent SARS-CoV-2-specific CD8+ T-cell memory is composed of heterogeneous subsets, e.g. T_CM_, T_EM1_, T_EM2_ and T_EM3_, required for a flexible response upon re-infection. Future studies have to evaluate whether differences in pre-existing and newly induced SARS-CoV-2-specific CD8+ T-cell responses are linked to different courses of infection. However, with this study, we now established the experimental tools for high resolution *ex vivo* analyses of SARS-CoV-2-specific CD8+ T cells required to answer the question about a potential rheostat action of virus-specific CD8+ T cells in SARS-CoV-2 pathogenesis versus protection.

## Acknowledgments

We thank all patients for participating in the current study and FREEZE-biobank-Center for biobanking of the Freiburg University Medical Center and the Medical Faculty for support. The study was funded by the Federal Ministry of Education and Research (Grant number 01KI1722) to GK, MH, MP, MS and RT and by a COVID-19 research grant of the Ministry of Science, Research and Art, State of Baden-Wuerttemberg to C.N.H. and B.B.. Work presented was also supported by the CRC/TRR 179-Project 01 and CRC 1160-Project A02 to R.T., CRC/TRR 179-Project 02 and CRC 1160-Project A06 to C.N.-H., CRC/TRR 179-Project 04 to T.B., CRC/TRR 179-Project 20 and CRC 1160-Project A02 to M.H., CRC/TRR 179-Project 21, CRC 1160-Project A03 and BE-5496/5-1 to B.B. of the German Research Foundation (DFG; TRR 179 project number: 272983813; CRC 1160 project number 256073931). M.H. was supported by a Margarete von Wrangell fellowship (State of Baden-Wuerttemberg). D.B. and T.B. are supported by the Berta-Ottenstein Programme, Faculty of Medicine, University of Freiburg. H.E.M. was supported by DFG ME3644/5-1 and D.A.P. by a Welcome Trust Senior Investigator Award (100326/Z/12/Z). The funding body had no role in the decision to write or submit the manuscript.

## Author contributions

I.S., J.K., V.O. and K.W. planned, performed and analyzed experiments with the help of S., F.D. and O.S. L.S. and S.K. performed and analyzed CyTOF data with the assistance of M. S. L.. I.S., J.K., V.O., K.W., L.S. and S.K. contributed equally to this work. A.D., H.L., B.Bi., D.B., S.R., C.F.W., A. N., D.D. were responsible for patient recruitment. H.M. and A.S. provided barcoding reagents for CyTOF analysis. S.L.-L. and D.P. provided influenza M1_58_/A*02 tetramers. F.E. performed four-digit HLA-typing by next generation sequencing. M.P. and D.H. performed antibody testing. M.S. and G.K. provided virological expertise and contributed to data interpretation. B.B. designed and supervised CyTOF analysis. T.B., R.T., M.H. and C.N.H. designed the study and contributed to experimental design and planning. I.S., J.K., V.O., R.T., M.H. and C.N.H. interpreted data and wrote the manuscript. C.N.H., M.H., R.T., B.B. and T.B. are shared last authors.

## Methods

### Study Cohort

A total of 26 COVID-19 convalescent individuals following a mild course of SARS-CoV-2 infection and 25 age and sex-matched historic controls (collected before August 2019) were recruited at the Freiburg University Medical Center, Germany. Mild course of infection was defined as clinical symptoms without signs of respiratory insufficiency. Patient characteristics are summarized in Extended Data Table I. SARS-CoV-2 infection was confirmed by positive PCR testing from oropharyngeal swab and/or SARS-CoV-2 spike IgG positive antibody testing in the presence of typical symptoms. Peptide-loaded major histocompatibility complex I (MHC I) tetramer-based magnetic bead enrichment of virus-specific CD8+ T cells was performed with samples from 18 SARS-CoV-2 convalescent individuals and 6 historic controls. HLA-typing was performed by next-generation sequencing. Influenza-specific CD8+ T-cell characterization was performed in 5 SARS-CoV-2 convalescent individuals. Written informed consent was obtained from all participants and the study was conducted according to federal guidelines, local ethics committee regulations (Albert-Ludwigs-Universität, Freiburg, Germany; vote #: 322/20) and the Declaration of Helsinki (1975).

### PBMC isolation

Venous blood samples were collected in EDTA-anticoagulated tubes. Peripheral blood mononuclear cells (PBMCs) were isolated with lymphocyte separation medium density gradients (Pancoll separation medium, PAN Biotech GmbH; Aidenbach, Germany) and resuspended in RPMI 1640 medium supplemented with 10% fetal calf serum, 1% penicillin/streptomycin, and 1.5% HEPES buffer 1 mol/L (complete medium; all additives from Thermo Scientific (Waltham, MA)) and stored at −80°C until used.

### Prediction of SARS-CoV-2-specific CD8+ T-cell epitopes

The entire viral amino acid sequence of SARS-CoV-2 (GenBank: MN908947.3) was analyzed for *in silico* peptide binding with ANN 4.0 on the Immune Epitope Database website^11^. The five best 8-,9- or 10-mer peptides calculated for the HLA alleles A*01:01, A*02:01, A*03:01, A*11:01, A*24:02, B*07:02, B*08:01, B*15:01, B*40:01, and B*44:02/03 were selected and synthesized for further analysis. Additionally, 13 epitopes that were predicted by Grifoni et al. with high sequence similarity to SARS-CoV-1 were included, summarized in Table 1^8^.

### Sequence Alignment

Sequence homology analyses were performed in Geneious Prime 2020.0.3 (https://www.geneious.com/) using Clustal Omega 1.2.2 alignment with default settings^12^. Reference genomes of human coronaviruses were downloaded from NCBI database 229E (NC_002645), HKU1 (NC_006577), NL63 (NC_005831), OC43 (NC_006213), MERS (NC_019843) and SARS-CoV-1 (NC_004718). Proteins of human coronaviruses were aligned according to their homology (amino acid level) only if the protein of interest has a homolog in the respective coronavirus. Confirmed SARS-CoV-2 epitopes were then mapped to the corresponding protein alignment, summarized in Extended Data Table II.

### Peptides and tetramers

Peptides were synthesized with an unmodified N-terminus and an amidated C-terminus with standard Fmoc chemistry and a purity of >70% (Genaxxon Bioscience, Ulm, Germany). HLA class I easYmers^®^ (immunAware, Copenhagen, Denmark) were loaded with peptide according to manufacturer’s instructions (A*01/ORF3a_207-215_, A*01/ORF1ab_4163-4172_, A*02/ORF3a_139-147_, B*07/N_105-113_) or ordered as peptide-loaded monomers (B*44:03/N_322-330_, B*44:03/ORF1ab_3946-3954_). SARS-CoV-2 peptide-loaded HLA class I tetramers were generated by conjugation of biotinylated peptide-loaded HLA class I easYmers^®^ with phycoerythrin (PE)-conjugated streptavidin (Agilent, Santa Clara, US) according to the manufacturer’s instructions. Influenza-specific HLA-A*02/M1_58-66_ (GILGFVFTL) tetramers were generated as described previously^13^.

### In vitro expansion of virus-specific CD8+ T-cells and assessment of effector function

PBMCs (1−2×10^6^) were stimulated with epitope-specific peptides (5 μM) and anti-CD28 mAb (0.5 μg/mL, BD) and expanded for 14 days in complete RPMI culture medium containing rIL2 (20 IU/mL, Miltenyi Biotec). The expansion factor was calculated based on peptide-loaded HLA class I tetramer staining as described before^14^. Cytokine production and degranulation were assessed 5 hours after restimulation with epitope-specific peptides as previously described^14^.

### Magnetic bead-based enrichment of antigen-specific CD8 T cells

Enrichment of virus-specific CD8+ T cells was performed as described before^15^. Briefly, 2-3×10^7^ peripheral blood mononuclear cells (PBMCs) were labelled for 30 min with PE-coupled peptide-loaded HLA class I tetramers. Subsequent enrichment was performed with anti-PE beads using MACS technology (Miltenyi Biotec, Germany) according to the manufacturer’s protocol. Enriched SARS-CoV-2-specific CD8+ T cells were used for multiparametric flow cytometry analysis. Frequencies of virus-specific CD8+ T cells were calculated as described previously^15^.

### Multiparametric flow cytometry

The following antibodies were used for multiparametric flow cytometry: anti-CCR7-PE-CF594 (150503, 1:50), anti-CCR7-BUV395 (3D12, 1:50), anti-CCR7-BV421 (150503, 1:33), anti-CD4-BV786 (L200, 1:200), anti-CD8-BUV496 (SK1, 1:100), anti-CD8-BUV510 (SK1, 1:100), anti-CD8-APC (SK-1, 1:200), anti-CD27-BV605 (L128, 1:200), anti-CD28-BV421 (CD28.2, 1:100), anti-CD28-BV711 (CD28.2, 1:100), anti-CD45RA-BV786 (HI100, 1:800), anti-CD45RA-BUV737 (HI100, 1:200), anti-CD69-BUV395 (FN50, 1:50), anti-CD107a-APC (H4A3, 1:100), anti-CD127-BV510 (HIL-7R-M21, 1:25), anti-EOMES-PerCP-eF710 (WD1928, 1:50), anti-IFN-γ-FITC (25723.11, 1:8), anti-IL-21-PE (3A3-N2.1, 1:25), anti-PD-1-BV786 (EH12.1, 1:33), anti-TNF-PE-Cy7 (Mab11, 1:400) (BD Biosciences, Germany). Anti-BCL-2-BV421 (100, 1:200), anti-CD25-BV650 (BC96, 1:33), anti-CD38-BV650 (HB-7, 1:400), anti-CD57-BV605 (QA17A04, 1:100), anti-CX3CR1-APC-eFluor660 (2A9-1, 1:50), anti-CXCR3-PerCP-Cy5.5 (G025H7, 1:33), anti-IL-2-PerCP-Cy5.5 (MQ1-17H12, 1:100), anti-IL17A-BV605 (BL168, 1:100), anti-PD-1-PE-Cy7 (EH12.2H7, 1:200), anti-rabbit-PE-CF594 (Poly4064, 1:200) anti-CD45RA-BV510 (HI100, 1:200), (BioLegend, UK), anti-FOXO1-pure (C29H4, 1:33), anti-TCF1-AlexaFluor488 (C63D9, 1:100) (Cell Signaling, Germany), anti-CD14-APC-eFluor780 (61D3, 1:400), anti-CD19-APC-eFluor780 (HIB19, 1:400), anti-CD27-FITC (0323, 1:100), anti-KLRG1-BV711 (13F12F2, 1:50), anti-T-BET-PE-Cy7 (4B10, 1:200), anti-TOX-eFluor660 (TRX10, 1:100) (eBioscience, Germany). A fixable Viability Dye (APC-eFluor780 1:200, 1:400) (eBioscience, Germany) or ViaProbe (7-AAD, 1:33) (BD Biosciences, Germany)) was used for live/dead discrimination. FoxP3/Transcription Factor Staining Buffer Set (eBioscience, Germany) and Fixation/Permeabilization Solution Kit (BD Biosciences, Germany) were applied according to the manufacturer’s instructions to stain for intranuclear and cytoplasmic molecules, respectively. Fixation of cells in 2% paraformaldehyde (PFA, Sigma, Germany) was followed by subsequent analyses on FACSCanto II, LSRFortessa (BD, Germany) or CytoFLEX (Beckman Coulter). Data analyses were performed with FlowJo 10 (Treestar, USA).

### Dimensionality reduction of multiparametric flow cytometry data

The visualization of multiparametric flow cytometry data was done with R version 4.0.2 using the Bioconductor (version: Release (3.11)) CATALYST package (Crowell H, Zanotelli V, Chevrier S, Robinson M (2020). CATALYST: Cytometry dATa anALYSis Tools. R package version 1.12.2, https://github.com/HelenaLC/CATALYST). The analyses were performed on gated virus-specific CD8+ T cells for two panels separately. Analysis of panel 1 (transcription factors) included the markers CD45RA, CCR7, CD27, CD28, BCL-2, TCF-1, CD69, CD38, PD-1, EOMES, T-BET and TOX. Analysis of panel 2 (surface markers) was performed on CCR7, CD45RA, CD27, CD28, CD25, CD127, CD57, KLRG1, CXCR3, PD-1, CX_3_CR1 and FOXO-1. Down sampling of cells to the number of cells present in the sample with the fewest cells was performed prior to dimensionality reduction in order to facilitate the visualization of different samples. Marker intensities were transformed by arcsinh (inverse hyperbolic sine) with a cofactor of 150. Dimensionality reduction on the transformed data was achieved by t-distributed stochastic neighbor embedding (t-SNE), multidimensional scaling (MDS) and Diffusion Map visualization.

### Mass cytometry

Mass cytometry reagents were obtained from Fluidigm or generated by custom conjugation to isotope-loaded polymers using MAXPAR X8 conjugation kit (Fluidigm). Mass cytometry antibodies used are shown in SI-Table 2. Mass cytometry tetramers were generated by tetramerization of pMHC monomers with Streptavidin conjugated to Eu^151^ using Lightning link conjugation kit (Expedon, Inc.) Sample barcoding was performed using anti-β2M barcodes, cells were then pooled and staining was performed as previously described^16^. Briefly, the single-cell suspension was pelleted, incubated with 20 μM Lanthanum-139 (Trace Sciences)-loaded maleimido-mono-amine-DOTA (Macrocyclics) in PBS for 10 min at RT for live/dead discrimination (LD). Cells were washed in staining buffer and resuspended in staining buffer containing tetramers, incubated for 30min at RT and washed twice. Cells were then resuspended in surface antibody cocktail, incubated for 30 min at RT, washed twice in staining buffer, pre-fixed with PFA 1.6%, washed, then fixed and permeabilized using FoxP3 staining buffer set (eBioscience) and stained intracellularly for 60 min at RT. Cells were further washed twice before fixation in 1.6% PFA (Electron Microscopy Sciences) solution containing 125 nM Iridium intercalator overnight at 4°C. Prior to data acquisition on a CyTOF Helios (Fluidigm), cells were washed twice in PBS and once in CAS. Mass cytometry data was analyzed after debarcoding and bead-based normalization. For analysis of mass cytometric data samples were first gated on Iridium intercalator positive, live, single CD45+CD3+CD8+ T cells using FlowJo (v10.6). CD8+ T cells were then exported for analysis in Omiq (Omiq, Inc.). Virus-specific CD8+ T cells were identified by manual gating. A workflow including dimension reduction using optSNE, PARC clustering analysis and Wishbone trajectory analysis was implemented in Omiq. Clustering and dimension reduction analysis were performed based on CD45RA, CD45RO, CCR7, CD28, CD127, CD16, CD25, CD26, CD38, CD39, CD56, CD57, CD69, CD103, CD161, CCR6, CCR9, CXCR3, CXCR5, CXCR6, CX3CR1, CRTH2, TCF-1, TOX, TIGIT, T-BET, EOMES, KLRG1 and PD-1. Further analysis and heatmap visualization was performed using R (v4.0) (https://www.r-project.org).

### SARS-CoV-2 spike IgG antibody determination

SARS-CoV-2 spike IgG antibodies were determined by the Euroimmune assay as described in the product instructions.

### Statistics

Statistical analysis was performed with GraphPad Prism 8 (USA). Statistical significance was assessed by Kruskal-Wallis testing including Dunn’s multiple comparisons test and Spearman correlation. (*P<0.05; **P<0.01; ***P<0.001; ****P<0.0001).

