## Supplementary Information for "*Ex vivo* detection of SARS-CoV-2-specific CD8+ T cells: rapid induction, prolonged contraction, and formation of functional memory"

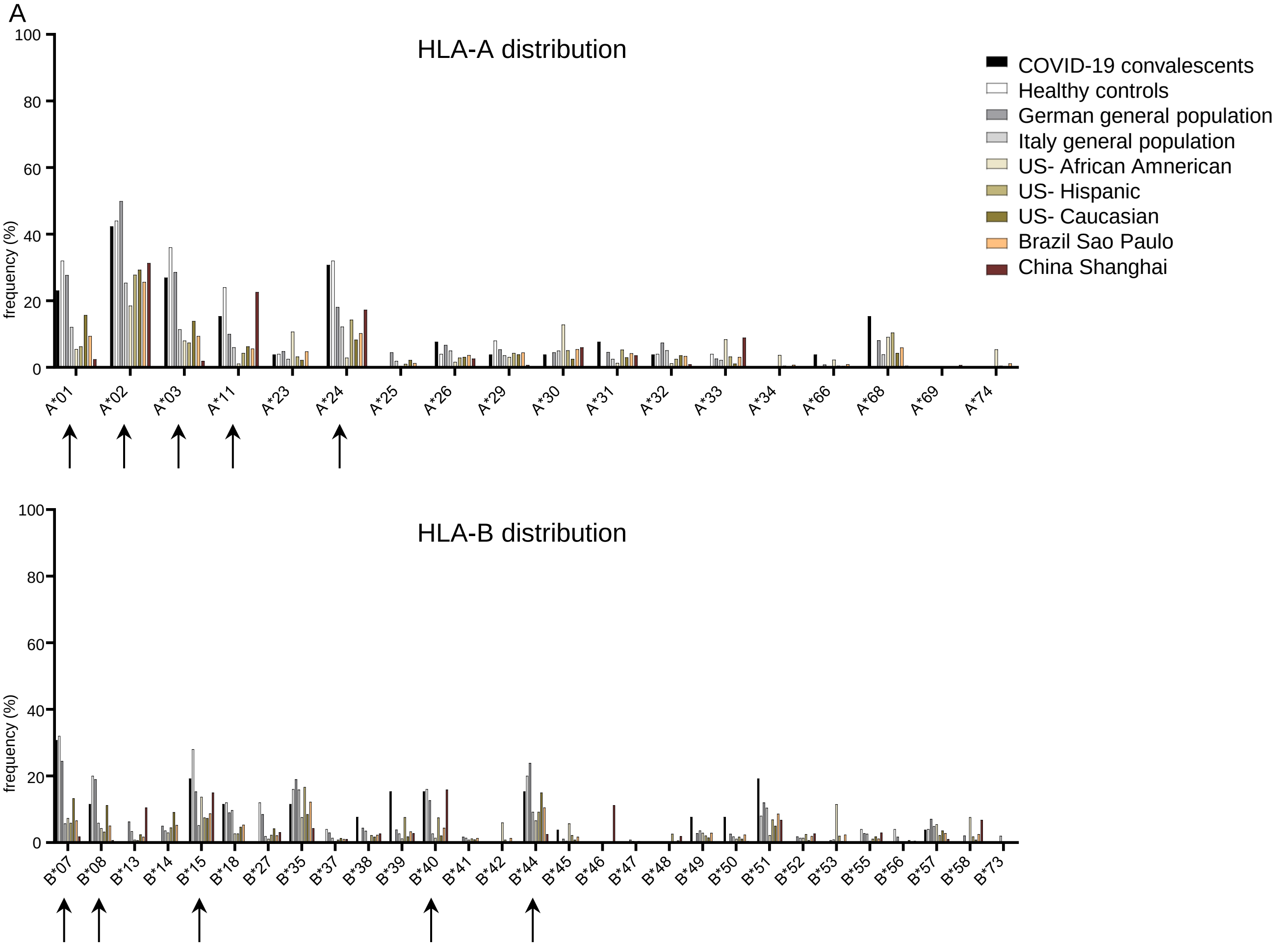


**Extended Data Figure 1:**
HLA-A and HLA-B distribution in different populations compared to the study population.
Arrows indicate HLA alleles for which peptide epitope candidates were predicted and further analyzed.

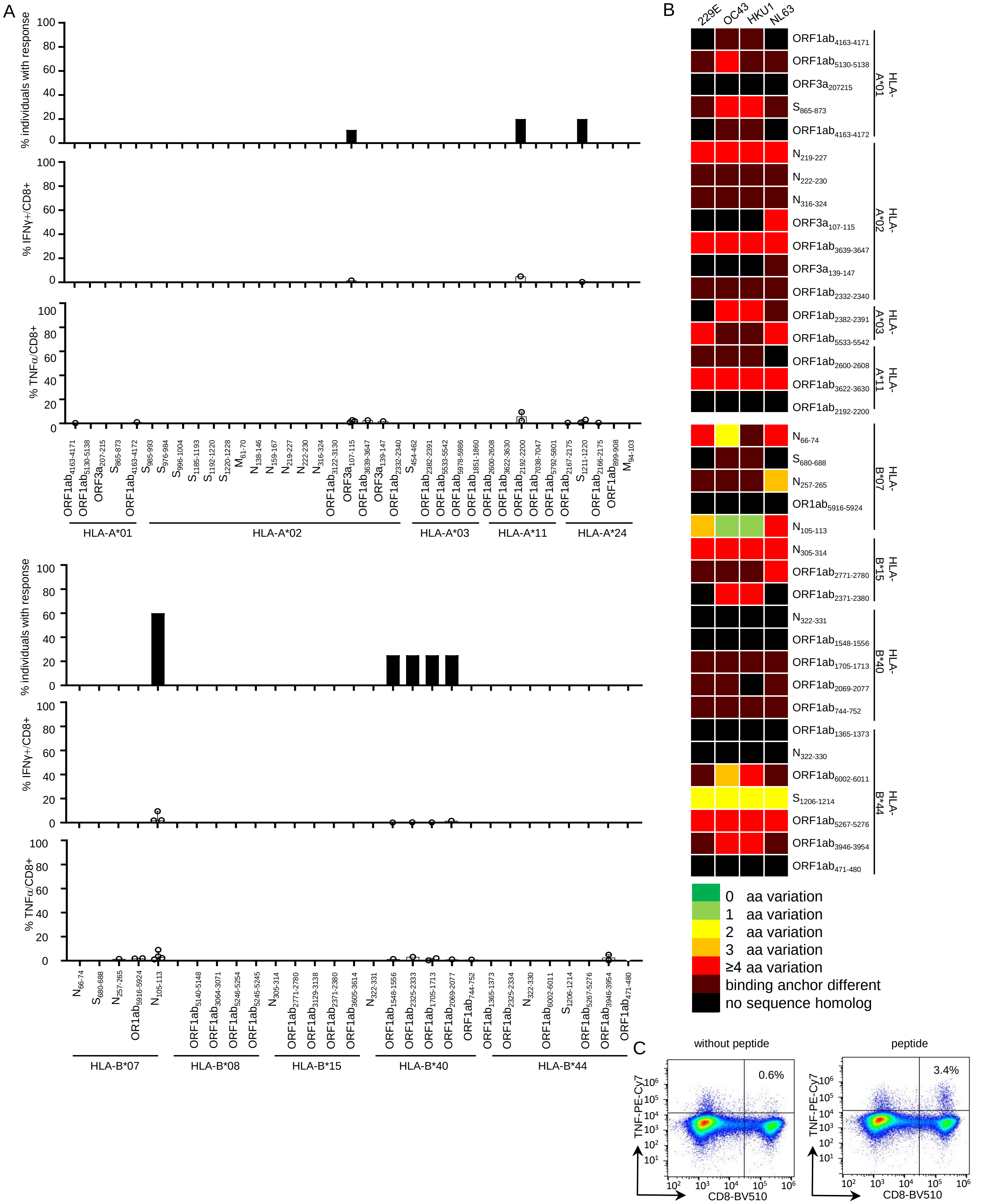


**Extended Data Figure 2:**

**(A)** % of historic controls with positive IFN-γ response towards HLA-A- and HLA-B-restricted SARS-CoV-2 peptides as well as the strength of individual responses as % IFN-γ+ and % TNF+ of CD8+ T cells. **(B)** Heat map illustrating the degree of homology between confirmed SARS-CoV-2 epitopes and "common cold" corona viruses 229E, OC43, HKU1 and NL63 (bright green: no amino acid (aa) change, 100% homology; light green: 1 aa difference; yellow: 2 aa difference; orange: 3 aa difference; bright red: 4-10 aa difference, dark red: aa residue at HLA-binding anchor is different, resulting in an IC50 at least ten times greater than predicted for the SARS-CoV-2 epitope, calculated by ANN 4.0 ; black: no homolog sequence). **(C)** Exemplary dot plot showing TNF production with and without SARS-CoV-2 peptide re-stimulation after 14-days *in vitro* expansion in a historic control. Bar charts show the median with IQR

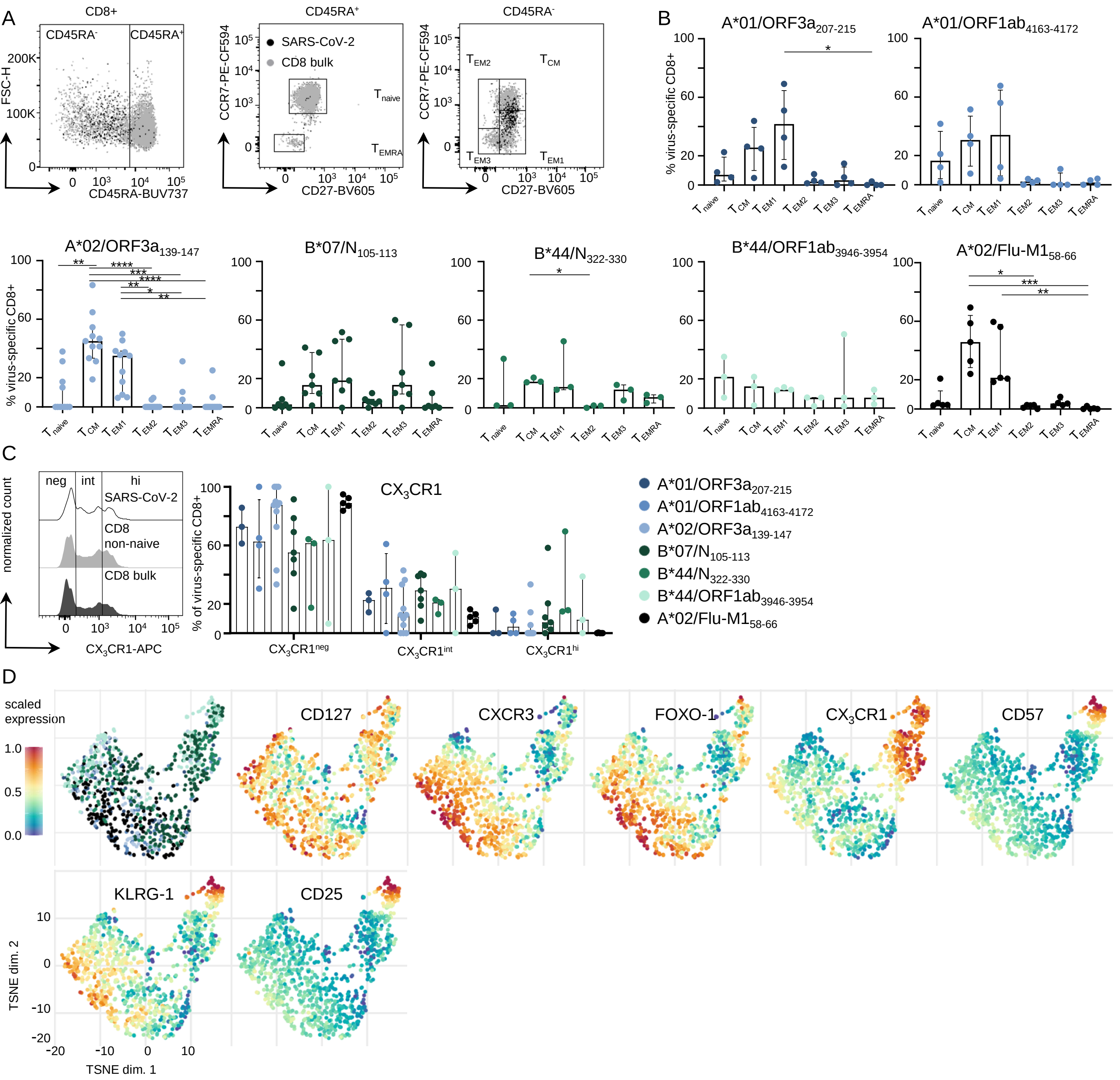


**Extended Data Figure 3:**

**(A)** Gating strategy for memory CD8+ T-cell population differentiation. **(B)** Distribution of CD8+ T-cell memory subsets, naïve T cells ($T_{naive}$), central memory T cells ($T_{CM}$), effector memory T cells 1 ($T_{EM1}$), effector memory T cells 2 ($T_{EM2}$), effector memory T cells 3 ($T_{EM3}$) and terminally differentiated effector memory cells re-expressing CD45RA ($T_{EMRA}$) between CD8+ T cells targeting the different epitopes. **(C)** Exemplary histogram showing the gating of cell populations expressing no (neg) $CX_3CR1$, intermediate (int) $CX_3CR1$ and high (hi) $CX_3CR1$ on CD8+ bulk (black) and non-naïve CD8+ T cells (grey) as well as SARS-CoV-2-specific CD8+ T cells (white). % of SARS-CoV-2-specific CD8+ T cells targeting the different epitopes expressing neg, int or hi levels of $CX_3CR1$. **(D)** t-SNE representation of flow cytometric data, which were derived from 18 convalescent SARS-CoV-2 individuals, comparing SARS-CoV-2-specific CD8+ T cells by their HLA restriction (left) and expression levels of CD127, CXCR3, FOXO-1, $CX_3CR1$, CD57, KLRG-1 and CD25 plotted on the t-SNE plot. Expression levels are color-coded: blue, low expression; red, high expressed. Bar charts show the median with IQR. Statistical significance was assessed by Kruskal-Wallis testing including Dunn's multiple comparisons test. (*P<0.05; **P<0.01; ***P<0.001; ****P<0.0001)

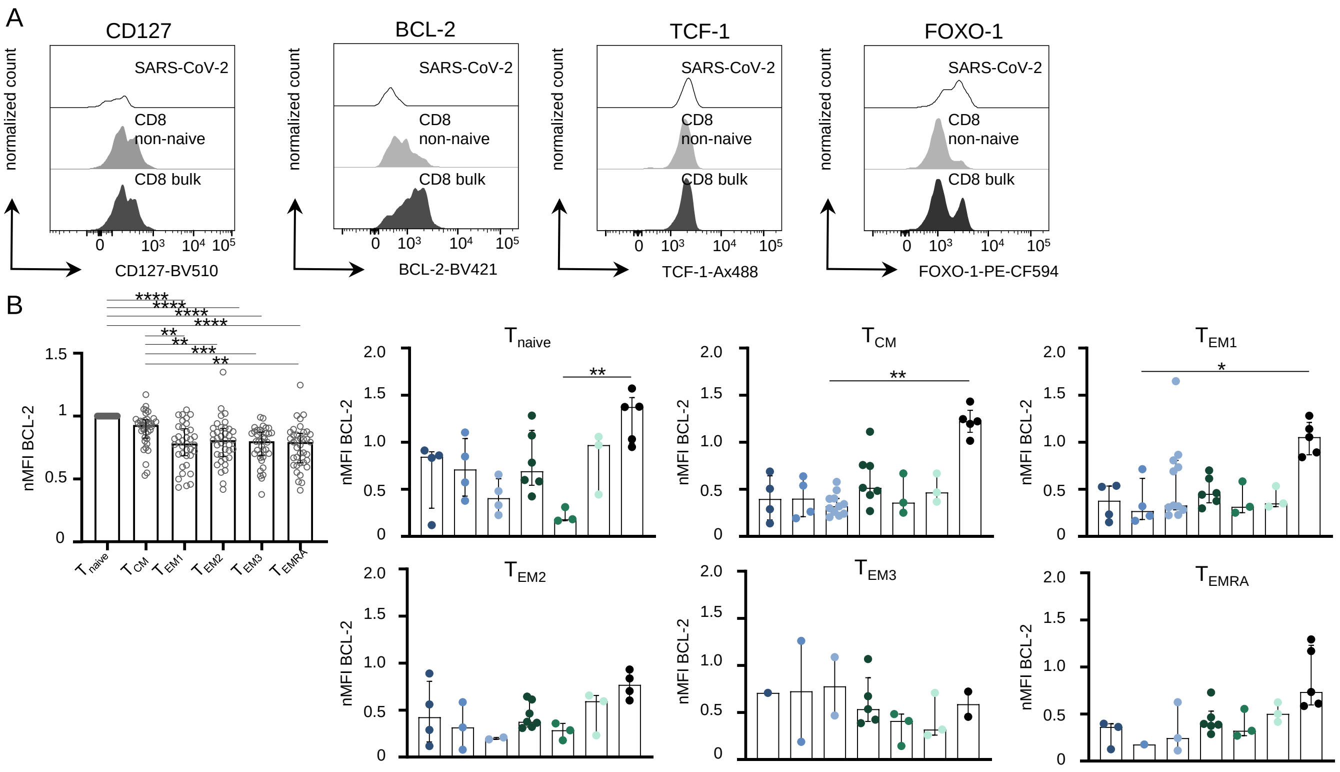


**Extended Data Figure 4:**

**(A)** Exemplary histograms depicting the expression levels of CD127, BCL-2, TCF-1 and FOXO-1 on CD8+ bulk (black) and non-naïve CD8+ T cells (grey) as well as SARS-CoV-2-specific CD8+ T cells (white). **(B)** BCL-2 expression of different memory cell populations on bulk CD8+ T cells (left) and of SARS-CoV-2-specific CD8+ T cells of the different memory CD8+ T-cell subsets. Bar charts show the median with IQR. Statistical significance was assessed by Kruskal-Wallis testing including Dunn's multiple comparisons test. (*P<0.05; **P<0.01; ***P<0.001; ****P<0.0001)

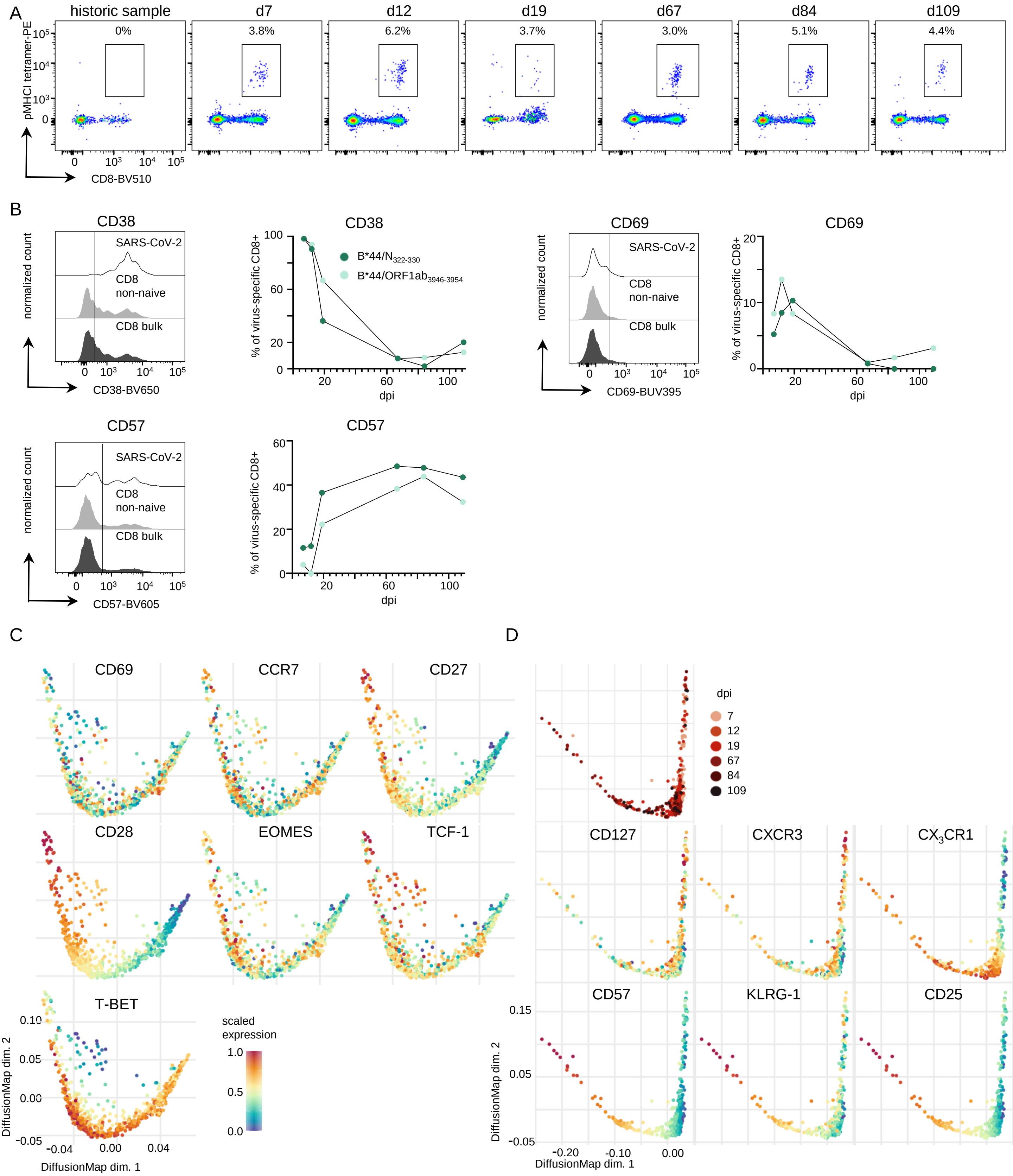


**Extended Data Figure 5:**
**(A)** Representative dot plots showing B*44/ORF1ab$_{3946-3954}$-specific CD8+ T cells *ex vivo* after tetramer-based enrichment. **(B)** Exemplary histograms depicting the protein expression levels on CD8+ bulk (black), non-naïve CD8+ T cells (grey) and SARS-CoV-2-specific CD8+ T cells (white) and the expression of these markers on SARS-CoV-2-specific CD8+ T cells days post infection (dpi). **(C)** Diffusion map representation of flow cytometric data, which were derived from longitudinal analysis from a convalescent SARS-CoV-2 individual, demonstrating the diffusion of B*44/N$_{322-330}$- and B*44/ORF1ab$_{3946-3954}$-specific T cells in relation to time post infection. Protein expression levels are plotted on the diffusion map. Expression levels are color-coded: blue, low expression; red, high expression. **(D)** Diffusion map representation of flow cytometric data, SARS-CoV-2-specific T cells in relation to time post infection which is distinguished by a color gradient from light (early time points) to dark red (late time points) color (top left). Protein expression levels are plotted on the diffusion map.

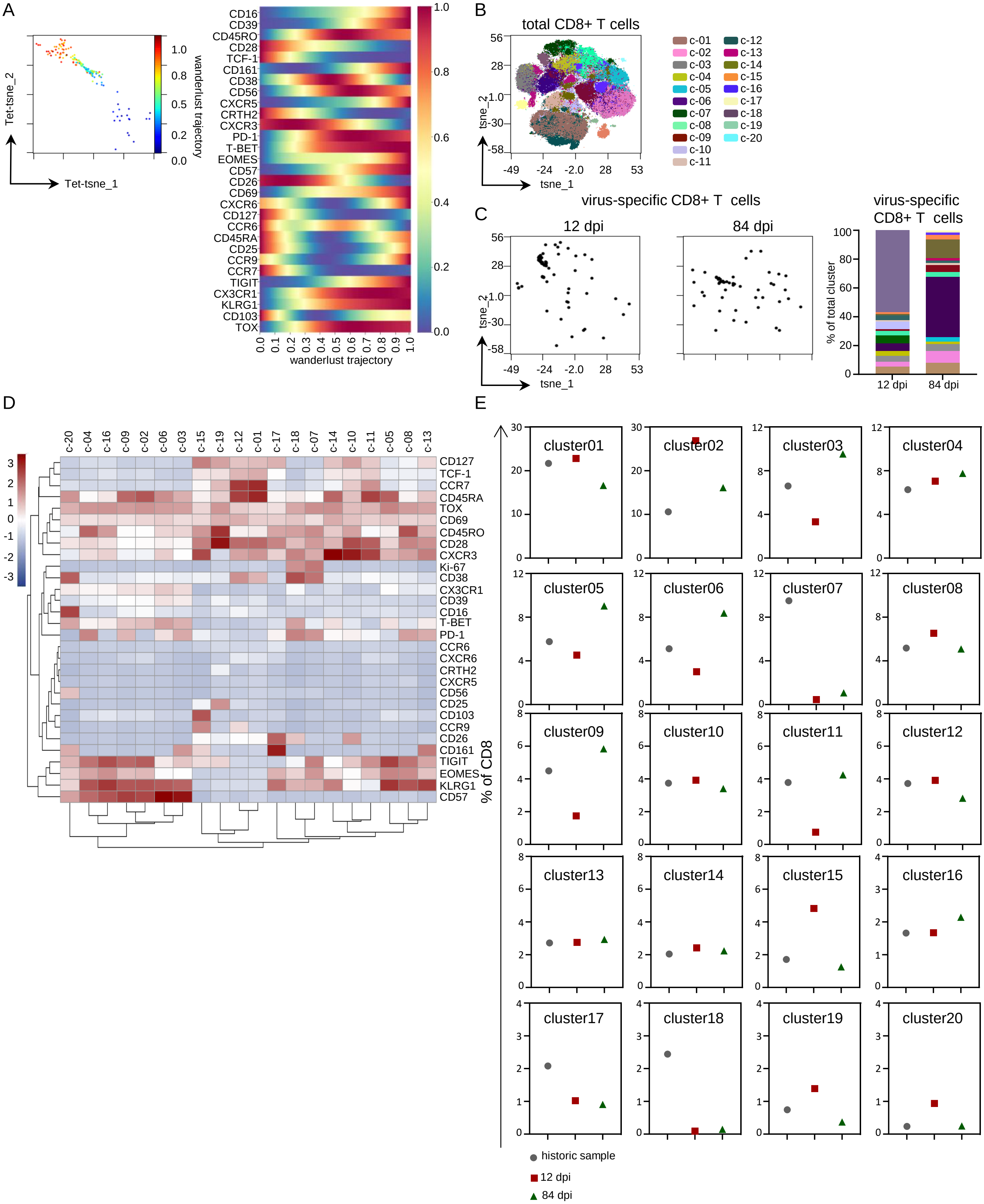


**Extended Data Figure 6:**
**(A)** tSNE map was calculated with all SARS-CoV-2-specific CD8+ T-cell data during acute and resolved infection (tet-SNE) and wanderlust trajectory analysis was performed on virus-specific CD8+ T cells and are shown colorized as indicated on the tet-SNE plot. Marker expression is depicted according to wanderlust trajectory progression. Expression levels are color-coded: blue, low expression; red, high expression. **(B)** The CD8 landscape in acute resolving SARS-CoV-2 was calculated using tSNE on CD8+ T cells from pre-infection, acute infection and resolved infection. Clustering was performed with PARC algorithm and clusters are indicated by the indicated color. **(C)** SARS-CoV-2-specific CD8+ T cells are displayed on the tSNE map. Frequency of virus-specific CD8+ T cells per CD8 cluster is illustrated using stacked bar chart. **(D)** Hierarchically clustered heatmap of PARC cluster phenotypes – the indicated marker expression is shown per cluster as z-Score of median signal intensity per channel; blue, low expression; red, high expression. **(E)** Frequency of each cluster of historic sample (circle), 12 dpi (square) and 84 dpi (triangle).

**Extended Data Table I:** Participants' characteristics

| | COVID-19 mild convalescents n=26 | Healthy controls n=25 |
|---|---|---|
| Male/female | 12/14 | 8/17 |
| Age *years: median (range)* | 32.5 (24-56) | 30 (24-78) |
| Sample collection date | March-June 2020 | May 2015-July 2019 |
| Days post onset of symptoms at sample collection (n=22) *median (range)* | 24 (14-70) | NA |
| SARS-CoV-2 Spike IgG positive | 77% (20/26) | 5% (1/20) |
| Symptoms | 85% (22/26) | NA |
| Cough | 54% (14/26) | NA |
| Headaches | 50% (13/26) | NA |
| Anosmia | 46% (12/26) | NA |
| Ageusia | 46% (12/26) | NA |
| Fever | 42% (11/26) | NA |
| Rhinorrhea | 31% (8/26) | NA |
| Sore throat | 23% (6/26) | NA |
| Myalgia | 19% (5/26) | NA |
| Dyspnea | 15% (4/26) | NA |
| Diarrhea | 15% (4/26) | NA |
| Fatigue | 8% (2/26) | NA |

**Extended Data Table II:** Sequence homology analyses between SARS-COV-2 epitopes and human coronaviruses

| **ORF1ab$_{471-480}$** | **EEIAIILASF** |
|---|---|
| SARS-CoV1 | ..V....... |
| MERS | DVVLA..SGT |
| 229E | SNVRRQ---- |
| OC43 | K.TNL.---- |
| HKU1 | K.T.L.---- |
| NL63 | DVKFAA---- |

| **ORF1ab$_{744-752}$** | **GETLPTEVL** |
|---|---|
| SARS-CoV1 | .DSHD.VLT |
| MERS | -------.S |
| 229E | LFPHNDRIK |
| OC43 | .SGSDFSLA |
| HKU1 | .VAESVI.E |
| NL63 | .VSK.NAID |

| **ORF1ab$_{1365-1373}$** | **QEILGTVSW** |
|---|---|
| SARS-CoV1 | E........ |
| MERS | --------- |
| 229E | D-------- |
| OC43 | -------G. |
| HKU1 | -------D. |
| NL63 | --------- |

| **ORF1ab$_{1548-1556}$** | **GEVITFDN---L** |
|---|---|
| SARS-CoV1 | ...LSL.K---. |
| MERS | ---LSA-CRA-- |
| 229E | LVLSSLTCNVSF |
| OC43 | .IFNKA-----T |
| HKU1 | .SFYKA-----T |
| NL63 | SN.MDV------ |

| **ORF1ab$_{1705-1713}$** | **GEAANFCAL** |
|---|---|
| SARS-CoV1 | .D....... |
| MERS | .DSTD.I.. |
| 229E | .DVEI.V.F |
| OC43 | .RP.R.V.. |
| HKU1 | .RPHRLV.. |
| NL63 | .DVGP.VSF |

| **ORF1ab$_{2069-2077}$** | **TEVVGDIIL** |
|---|---|
| SARS-CoV1 | .....NV.. |
| MERS | PF.KDNVSF |
| 229E | SG.AYTAFS |
| OC43 | FK.EDSV.V |
| HKU1 | ----DA..V |
| NL63 | .I.SEK.SV |

| **ORF1ab$_{2192-2200}$** | **ASMPTTIAK** |
|---|---|
| SARS-CoV1 | ..L...... |
| MERS | .L---KT.G |
| 229E | .KA.QRT-- |
| OC43 | YT--.E..S |
| HKU1 | YT--.E..S |
| NL63 | .KA.KRTGV |

| **ORF1ab$_{2332-2340}$** | **ILFTRFFYV** |
|---|---|
| SARS-CoV1 | M...K...L |
| MERS | M.Y.SA.NW |
| 229E | ----LLYF. |
| OC43 | A.Y.AW..P |
| HKU1 | S.Y.VW..P |
| NL63 | LI.GNMYLR |

| **ORF1ab$_{2371-2380}$** | **LVQMAPISAM** |
|---|---|
| SARS-CoV1 | I.....V... |
| MERS | -----.MAGL |
| 229E | D.ICDE---- |
| OC43 | .AN.L.AHVF |
| HKU1 | VAN.L.AFVL |
| NL63 | -----.FDVL |

| **ORF1ab$_{2382-2391}$** | **RMYIFFASFY** |
|---|---|
| SARS-CoV1 | .......... |
| MERS | ...NLL.CLW |
| 229E | --LLVTVIVI |
| OC43 | .F..II...I |
| HKU1 | .F..VVTAM. |
| NL63 | NEFLATFIVC |

| **ORF1ab$_{2600-2608}$** | **STFNVPMEK** |
|---|---|
| SARS-CoV1 | A..S..... |
| MERS | .LY..TRD. |
| 229E | NS.GKDLNA |
| OC43 | .M.D.DKKS |
| HKU1 | .H.D.DRKS |
| NL63 | NS.FK---- |

| **ORF1ab$_{2771-2780}$** | **KQLIKVTLVF** |
|---|---|
| SARS-CoV1 | .LML.A..LC |
| MERS | .GYVLA.IIV |
| 229E | WLWLLCG..C |
| OC43 | YVCFVLS..C |
| HKU1 | YI.FF.S.IC |
| NL63 | YVCLF.VAL. |

| **ORF1ab$_{3622-3630}$** | **SAFAMMFVK** |
|---|---|
| SARS-CoV1 | A.C..LL.. |
| MERS | M..V.LL.. |
| 229E | .LCLTFV.. |
| OC43 | ISL..LL.. |
| HKU1 | VS.M.LL.. |
| NL63 | .LVLT..L. |

| **ORF1ab$_{3639-3647}$** | **FLLPSLATV** |
|---|---|
| SARS-CoV1 | ......... |
| MERS | ....VAICL |
| 229E | .....IIVA |
| OC43 | YIT.V.F.L |
| HKU1 | YII.V.C.L |
| NL63 | ....TVIAT |

| **ORF1ab$_{3946-3954}$** | **SEFSSLPSY** |
|---|---|
| SARS-CoV1 | ......... |
| MERS | ....H.ATF |
| 229E | .S.VGM..F |
| OC43 | ...VNMA.F |
| HKU1 | ...VNMA.F |
| NL63 | .S.V.M... |

| **ORF1ab$_{4163-4171}$** | **CTDDNALAY** |
|---|---|
| SARS-CoV1 | ......... |
| MERS | .NT-SS... |
| 229E | TS--EGN.L |
| OC43 | .NT-PTQC. |
| HKU1 | .NI-PTQC. |
| NL63 | LG--DGN.L |

| **ORF1ab$_{4163-4172}$** | **CTDDNALAYY** |
|---|---|
| SARS-CoV1 | .......... |
| MERS | .NT-SS.... |
| 229E | TS--EGN.L. |
| OC43 | .NT-PTQC.. |
| HKU1 | .NI-PTQC.. |
| NL63 | LG--DGN.L. |

| **ORF1ab$_{5130-5138}$** | **DTDFVNEFY** |
|---|---|
| SARS-CoV1 | .HE..D... |
| MERS | .PK..DKY. |
| 229E | .ES..DD.. |
| OC43 | .ST..T.Y. |
| HKU1 | .YT....Y. |
| NL63 | EES.IDDY. |

| **ORF1ab$_{5267-5276}$** | **QEYADVFHLY** |
|---|---|
| SARS-CoV1 | .......... |
| MERS | I..QN..WV. |
| 229E | P..RK..YAL |
| OC43 | E..QK..RV. |
| HKU1 | E..QK..RV. |
| NL63 | S..RK..YVL |

| **ORF1ab$_{5533-5542}$** | **VVYRGTTTYK** |
|---|---|
| SARS-CoV1 | .......... |
| MERS | .S.KSS.... |
| 229E | .T.KS.A.T. |
| OC43 | .Y..A..... |
| HKU1 | .Y..A..... |
| NL63 | .T.KS.V.T. |

| **ORF1ab$_{5916-5924}$** | **IPRRNVATL** |
|---|---|
| SARS-CoV1 | ......... |
| MERS | FTNYK---. |
| 229E | MTD-----. |
| OC43 | LDKVPQVET |
| HKU1 | LDKIQNTLP |
| NL63 | HAD-----. |

| **ORF1ab$_{6002-6011}$** | **EEAIRHVRAW** |
|---|---|
| SARS-CoV1 | .......... |
| MERS | ...V.Q..S. |
| 229E | DF.M....G. |
| OC43 | ...VKR.... |
| HKU1 | D...KR..G. |
| NL63 | DF...N..G. |

| **S$_{680-688}$** | **SPRRARSVA** |
|---|---|
| SARS-CoV1 | --SLL..TS |
| MERS | T..SV-RSV |
| 229E | --AV----- |
| OC43 | --..S.GAI |
| HKU1 | .S..K.RSI |
| NL63 | --PV----- |

| **S$_{865-873}$** | **LTDEMIAQY** |
|---|---|
| SARS-CoV1 | ...D...A. |
| MERS | MDVN.E.A. |
| 229E | ADA.RM.M. |
| OC43 | .SENQ.SG. |
| HKU1 | .SESQ.SG. |
| NL63 | ADA.RM.M. |

| **S$_{1206-1214}$** | **YEQYIKWPW** |
|---|---|
| SARS-CoV1 | ......... |
| MERS | .TY.N.... |
| 229E | V.T...... |
| OC43 | ..Y.V.... |
| HKU1 | ..M.V.... |
| NL63 | F.N...... |

| **ORF3a$_{107-115}$** | **YLYALVYF---L** |
|---|---|
| SARS-CoV1 | .....I.---. |
| MERS | LVFS.S-LL-VT |
| NL63 | ...KNFS.V-LF |

| **ORF3a$_{139-147}$** | **LLYDANYFL** |
|---|---|
| SARS-CoV1 | ........V |
| MERS | --------- |
| NL63 | SF.ENRFAA |

| **ORF3a$_{207-215}$** | **FTSDYYQLY** |
|---|---|
| SARS-CoV1 | ..EV....E |
| MERS | .DVG.S--- |
| NL63 | NGAVI.IV. |

| **N$_{66-74}$** | **FPRGQGVPI** |
|---|---|
| SARS-CoV1 | ......... |
| MERS | ..P.....L |
| 229E | VIPRNL... |
| OC43 | .VE...... |
| HKU1 | .SD...... |
| NL63 | VIPRNL... |

| **N$_{105-113}$** | **SPRWYFYYL** |
|---|---|
| SARS-CoV1 | ......... |
| MERS | A.......T |
| 229E | ..KLH.... |
| OC43 | L........ |
| HKU1 | L........ |
| NL63 | P.KVH.... |

| **N$_{219-227}$** | **LALLLLDRL** |
|---|---|
| SARS-CoV1 | ......... |
| MERS | GD..Y..L. |
| 229E | .KS.GF.KP |
| OC43 | I.S.V.AK. |
| HKU1 | I.N.V.AK. |
| NL63 | .KN.GF.NQ |

| **N$_{222-230}$** | **LLLDRLNQL** |
|---|---|
| SARS-CoV1 | ......... |
| MERS | .Y..L..R. |
| 229E | .GF.KPQEK |
| OC43 | .V.AK.GKD |
| HKU1 | .V.AK.GKD |
| NL63 | .GF.NQSKS |

| **N$_{257-265}$** | **KPRQKRTAT** |
|---|---|
| SARS-CoV1 | ........ |
| MERS | .M.H...S. |
| 229E | ...W..QPN |
| OC43 | ......SPN |
| HKU1 | .......PN |
| NL63 | ...W..VP. |

| **N$_{305-314}$** | **AQFAPSASAF** |
|---|---|
| SARS-CoV1 | .......... |
| MERS | .EL..T.... |
| 229E | .ELV..TA.M |
| OC43 | .EL..T.G.. |
| HKU1 | .EL..TPG.. |
| NL63 | .ELI.NQA.L |

| **N$_{316-324}$** | **GMSRIGMEV** |
|---|---|
| SARS-CoV1 | ......... |
| MERS | ...QFKLTH |
| 229E | FD.H.VSKE |
| OC43 | FG..LELAK |
| HKU1 | FG.KLDLVK |
| NL63 | FD.EVSTDE |

| **N$_{322-330}$** | **MEVT-----------PSGTW** |
|---|---|
| SARS-CoV1 | ....-----------..... |
| 229E | SKES-----------GNTVV |
| MERS | LTHQ-----NNDDHGNPVYF |
| OC43 | LAKVQNLSGNPDEPQKDVYE |
| NL63 | TDEV-----------GDNVQ |
| HKU1 | IDTAGVL------------- |

| **N$_{322-331}$** | **MEVT-----------PSGTWL** |
|---|---|
| SARS-CoV1 | ....-----------...... |
| MERS | LTHQ-----NNDDHGNPVYF. |
| 229E | SKES-----------GNTVV. |
| OC43 | LAKVQNLSGNPDEPQKDVYE. |
| HKU1 | IDTAGVL-------------- |
| NL63 | TDEV-----------GDNVQI |

**Extended Data Table III**: Mass cytometry panel

| Channel | Antigen | Clone |
|---|---|---|
| 89Y | CD45 | HI30 |
| 102-110 Pd | ß2m-Barcode | |
| 111 Cd | CD4 | RPA-T4 |
| 112 Cd | CD3 | UCHT1 |
| 113 In | CD39 | A1 |
| 114 Cd | CD45 RO | UCHL1 |
| 115 In | CD57 | TB01 |
| 116 Cd | CD8 | RPA-T8 |
| 139 La | MM-DOTA | |
| 140 etc | Beads | |
| 141 Pr | CCR6 | G034E3 |
| 142 Nd | CD26 | BA5b |
| 143 Nd | Nkp46 | 9,00E+02 |
| 144 Nd | CD69 | FN50 |
| 145 Nd | CD19 | HIB19 |
| 146 Nd | Ki-67 | B56 |
| 147 Sm | CD45RA | H100 |
| 148 Nd | CXCR6 | K041E5 |
| 149 Sm | CD25 | 2A3 |
| 150 Nd | CD127 | HIL-7R-M21 |
| 151 Eu | PE (Tetramer) | |
| 152 Sm | CD21 | BL13 |
| 153 Eu | TCR Va7.2 | 3C10 |
| 154 Sm | CCR9 | L053E8 |
| 155 Gd | CRTH2 | BM16 |
| 156 Gd | CXCR3 | G025H7 |
| 158 Gd | PD-1 | EH12.2H7 |
| 159 Tb | CCR7 | G043H7 |
| 160 Gd | Tbet | 4B10 |
| 161 Dy | CD28 | CD28.2 |
| 162 Dy | FoxP3 | PCH101 |
| 163 Dy | TCF-1 | C398.4A |
| 164 Dy | CD161 | HP3G10 |
| 165 Ho | EOMES | WD1928 |
| 167 Er | CD38 | HIT2 |
| 169 Tm | TIGIT | MBSA43 |
| 170 Er | CXCR5 | RF8B2 |
| 171 Yb | TCRgd | B1 |
| 172 Yb | CX3CR1 | K0124E1 |
| 173 Yb | KLRG1 | 13F12 |
| 174 Yb | CD103 | Ber-ACT8 |
| 175 Lu | CCR4 | L291H4 |
| 176 Yb | CD56 | NCAM16.2 |
| 191/193 | Iridium | |
| 198 Pt | ß2m-Barcode | |
| 209 Bi | CD16 | 3G8 |
